# A multi-organ map of the human immune system across age, sex and ethnicity

**DOI:** 10.1101/2023.06.08.542671

**Authors:** S Mangiola, M Milton, N Ranathunga, CSN Li-Wai-Suen, A Odainic, E Yang, W Hutchison, A Garnham, J Iskander, B Pal, V Yadav, JFJ Rossello, VJ Carey, M Morgan, S Bedoui, A Kallies, AT Papenfuss

## Abstract

Understanding tissue biology’s heterogeneity is crucial for advancing precision medicine. Despite the centrality of the immune system in tissue homeostasis, a detailed and comprehensive map of immune cell distribution and interactions across human tissues and demographics remains elusive. To fill this gap, we harmonised data from 12,981 single-cell RNA sequencing samples and curated 29 million cells from 45 anatomical sites to create a comprehensive compositional and transcriptional healthy map of the healthy immune system. We used this resource and a novel multilevel modelling approach to track immune ageing and test differences across sex and ethnicity. We uncovered conserved and tissue-specific immune-ageing programs, resolved sex-dependent differential ageing and identified ethnic diversity in clinically critical immune checkpoints. This study provides a quantitative baseline of the immune system, facilitating advances in precision medicine. By sharing our immune map, we hope to catalyse further breakthroughs in cancer, infectious disease, immunology and precision medicine.

## Introduction

Understanding the cellular and molecular heterogeneity across tissues in the human population is crucial to improving precision medical research. For example, immunotherapy and immunodiagnosis^1^ research would greatly benefit from a quantification of the diversity of the immune system across tissues, age^2^, sex^3^, and ethnicity^4,5^ with high resolution and coverage to target conserved and universal checkpoints.

Previous multi-tissue analyses of single cells have been limited to <400,000 cells^6^, included only a few tissues^7^ or individuals^8^, and mainly focused on cell type classification and early development. For example, Domínguez Conde et al.^8^ used a balanced experimental design with multiple tissues probed across individuals, with the benefit of having more control over the inter-subject variability. However, an analysis limited to an *ad hoc* cohort (e.g. n=12^8^) lacks the power to deconvolve the complexity of the human demographics.

Large-scale single-cell sequencing initiatives, such as the Human Cell Atlas^9^, have created an unprecedented opportunity to develop a comprehensive map that could yield invaluable insights into how immune cells are distributed across various tissues and how they function and communicate. Moreover, the number of studies using single-cell sequencing to understand immune cell functions is growing rapidly, providing a continuous stream of data for comprehensive analyses. However, the disparate nature of the individual data sources, the use of different sequencing approaches, and variations in experimental designs across studies provide significant barriers to such integrative data analyses. For example, despite the large sample size, sparsity remains across some combinations of factors (e.g. sex, age range, ethnicity, technology, tissue); that is, the number of samples in each combination of factors is highly variable. Also, the components of complex datasets are often hierarchically organised (e.g. a database including many tissues, the data of which has been produced by many studies and contributed by many samples). Overcoming these significant complexities requires data harmonisation and novel methodologies for scalable analysis. Data representation biases and sparsity, and hierarchical data organisation, characteristics of large-scale resources, have been addressed by multilevel models^10,11^. However, current models are not tailored to compositional analyses of single-cell data.

Here, we used novel methodologies to characterise a multi-organ map of the human immune system generated by harmonising transcriptomic data from almost 29 million cells across 12,981 samples and 31 anatomical locations into a single comprehensive resource. We developed a novel multilevel strategy for compositional analysis and used established multilevel modelling methods for transcriptomics^12^ and cell-communication analyses^13^. In ageing, we detangled primary and secondary programs and conserved compositional trajectories conserved across tissues and unique for tissue clusters. We resolved the link between ageing and the loss of immune adaptability and disruption of central immune processes. Moreover, our analyses articulate a pronounced sexual dimorphism in ageing, governed by a stark dichotomy in sex-dependent differential ageing trajectories. In a landscape marked by significant ethnic immune diversity, our findings lead to the implications for investigating universal therapeutic targets, accentuating our clear picture of a European predominance in data representation.

This study provides a whole-body map of immune cell composition and transcriptional activity, spanning tissues and traits of the human population. This compendium stands as an extensive and quantitative touchstone of the human immune system in its homeostatic state, representing a multifaceted reference for immunological and precision medicine investigation.

## Results

### Cell-atlas harmonisation and cell-type reannotation allow a comprehensive investigation of the immune system

Mining highly heterogeneous data sources like the Human Cell Atlas^9^ requires extensive data harmonisation (Figure 1). To this end, we standardised genes and sample identifiers, cell metadata such as tissue labels and age, and transformed gene-transcript abundance to a common scale. This curation resulted in a comprehensive resource facilitating detailed interrogation of human biology at the single-cell level across developmental stages, from embryonic to geriatric, sexes, ethnic groups and a wide range of lymphoid and non-lymphoid tissues and anatomical compartments. We organised the analyses in four parts, focusing on (i) immune ageing, (ii) sex dimorphism, (iii) sex-differential ageing, and (iv) ethnicity. For each analysis stream, we estimated changes in immune cellularity (proportion of immune vs non-immune cells) and composition at the body and tissue level.

**Figure 1.**
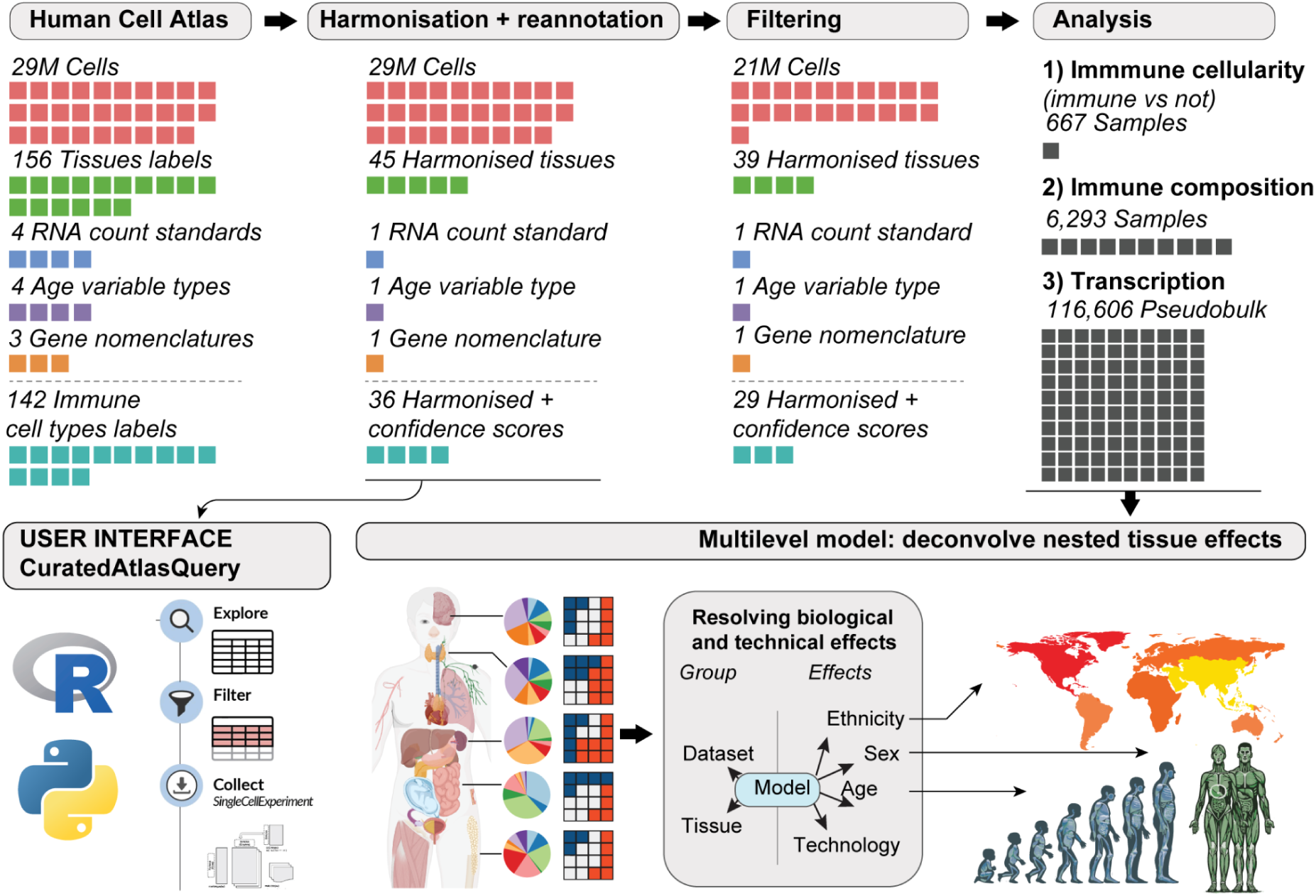
Overview of the study. Each coloured cell represents a quantity specified in its headers. From left to right: We retrieved Human Cell Atlas data through the CELLxGENE repository. We harmonised cell metadata, gene nomenclature, and count distributions and scored cell annotations for quality using an ensemble approach of several annotation references. We provide this resource to the community through the *CuratedAtlasQuery* interface. To improve analysis robustness, we filtered cells based on annotation quality and data abundance (e.g. excluding samples with less than 30 cells and studies with less than three samples). We analysed the association of immune cellularity (proportion of immune vs non-immune cells), composition and transcription with age, sex, and ethnicity (i.e. nine total analyses).

To enable the accurate, comprehensive investigation of the immune system, we also standardised cell-type annotations across studies and applied an ensemble annotation strategy to minimise the impact of study-specific standards (Figure 2A). We used the original annotations from each study and extended these using Azimuth PBMC^14^, Blueprint^15^ and Monaco^16^. Our ensemble approach revealed that 87% of cell identities had absolute consensus (among all four sources; Figure 2B). We created confidence tiers for cells with relative consensus (among most sources) or lack of consensus (Figure 2B, Methods). While confidence tier 1 is reserved for absolute consensus, confidence tiers 2 and 3 are reserved for relative consensus (among 3 and 2 sources, respectively), and 4 is reserved for lack of consensus. Notably, the proportion of cells assigned to each confidence tier varied across cell-profiling technologies (Figure 2C). While 10X-based datasets, Smart-seq2, Seq-well and micro-well reached a complete consensus for most cells, 10X ATAC-seq (used only as cell annotation; i.e. cell annotation), less than 25% of cells from Sci-seq, Slide-seq, and 10X Visium were in confidence tier 1. The proportion of cells in confidence tier 1 is also associated with ethnicity, tissue, and cell type. (Figure 2C, Supplementary Figure S1). Low-confidence annotations were not significantly associated with cell library size (Figure S2).

**Figure 2.**
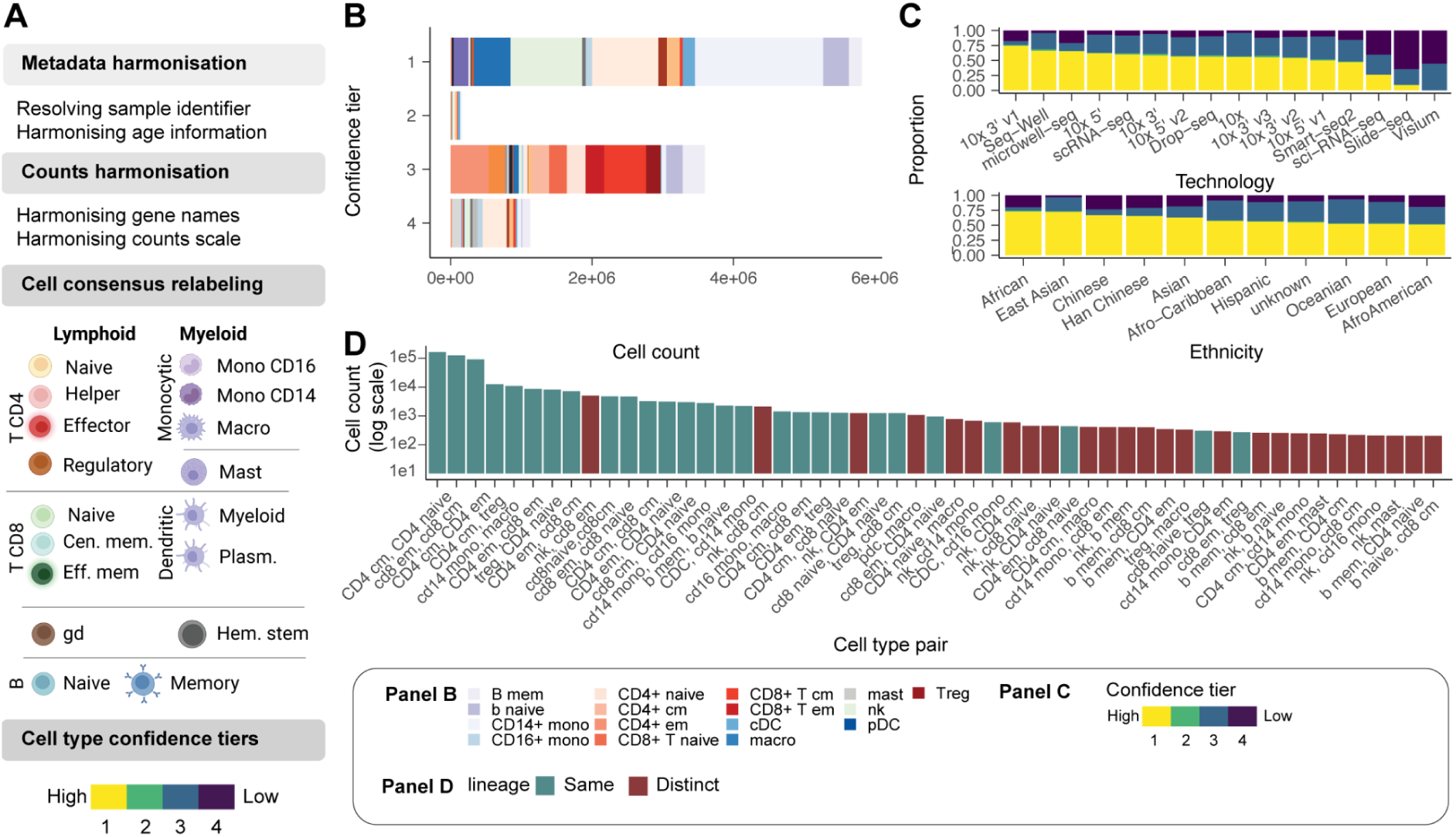
Data curation and query interface. A: Schematic of the curation pipeline. **B:** Number of immune cells in each confidence tier. **C:** Proportion of cells in each confidence tier across technologies and ethnicity. **D:** Co-occurrence of cell type annotations (between methods) for cells in confidence tiers 2-4. For example, Azimuth may annotate a cell as a CD4 T cell, while SingleR with Monaco reference may annotate it as a CD8 T cell.

We next sought to analyse the cell types for which the four annotations most often disagreed (Figure 2D). This analysis aimed to identify which cell types were most challenging to discern for the reference-based annotation approach. Most cell annotation disagreements represented activation states and subsets of common cell lineages (e.g., memory and naive CD4+ T cells). However, disagreement exists between more distantly related but transcriptionally convergent cell types such as NK and memory CD8+ T cells. Our harmonisation pipeline standardised a multi-study large-scale dataset across genes, samples, cells, and transcription quantifications, making it amenable to integrative analyses.

To support data access and facilitate systematic exploration of the harmonised and reannotated atlas, we developed a software interface, *CuratedAtlasQuery*. This enabled cells of interest to be selected based on their ontology and tissue of origin and incorporate key information about the age, sex, ethnicity and health status of the donor. For example, users can select CD4+ T cells across lymphoid tissues. The data for the selected cells can be downloaded locally into popular single-cell data containers^17^. Pseudobulk counts are also available to facilitate large-scale, summary analyses of transcriptional profiles. This platform offers a standardised workflow for accessing atlas-level datasets programmatically and reproducibly.

### Sccomp and tidybulk tools facilitate atlas-level compositional and transcriptional analyses

Gene expression atlases are often compiled from a mosaic of studies. This aggregation poses significant challenges to statistical analysis due to the absence of a uniform experimental design. The first layer of complexity is the tree structure of our Human Cell Atlas-derived resource. This resource amalgamates data across various tissues, each derived from multiple studies (Figure 3A). A notable proportion of these studies (26.2%) encompass multiple tissues.

**Figure 3.**
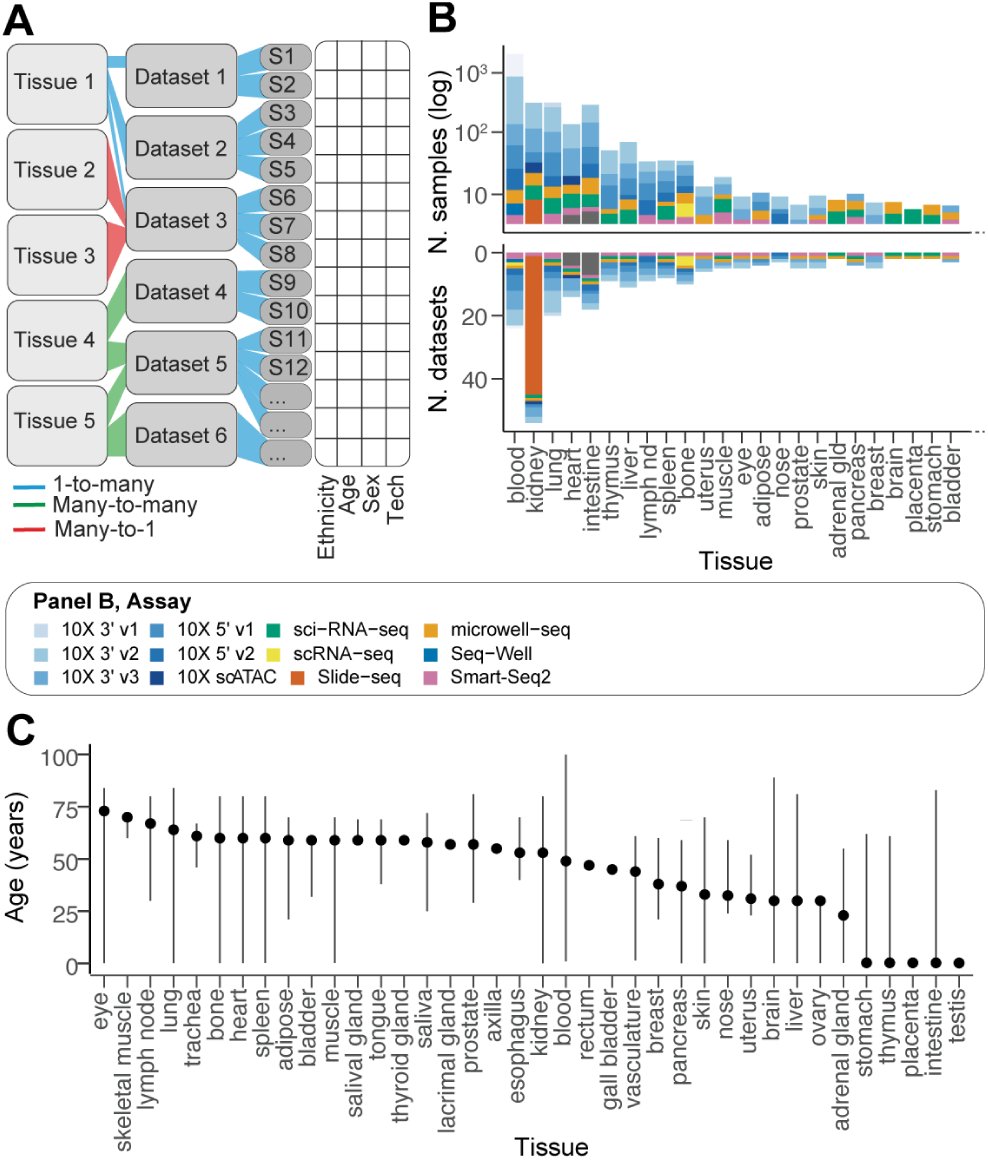
Complexity and biases of the data source. A: The schematic shows the multilevel structure of the data. Relationships between data strata are colour-coded. Tissues and datasets are used here as a hierarchical grouping. **B:** Summary statistics of the data content by tissue (using the harmonised tissue annotation). The bottom panel y-axis is log-scaled. The bars are coloured by sample processing technology. **C:** Age distribution across tissues. Lines define the range (from minimum to maximum). Points define the median.

A second layer of complexity is the data coverage bias. For example, there is a stark disparity in tissue representation, as evidenced by the two orders of magnitude difference between the most and least sampled tissues (Figure 3B). The blood samples are the most frequent, constituting 28% (3,582 samples) of the entire atlas. This tissue is closely followed by brain and heart samples, which account for 30% of the data. Conversely, tissues like the appendix vermiformis, gall bladder, and rectum are significantly under-represented, collectively comprising merely 0.02% of the atlas.

A third layer of complexity arises from the variable age distribution among tissue samples (Figure 3C). While blood, lung, and kidney samples exhibit a wide age range, others, such as adipose tissue and lymph nodes, are limited to a narrower age spectrum. These discrepancies underscore the intricacies of modelling cell and gene transcript abundance across diverse tissues and demographic variables.

This complexity is not suitable to standard single-level (i.e. fixed-effect) linear models, as the estimation of body-level effects (i.e. across all tissues) would be dominated by highly represented tissues (e.g. blood), and continuous effects like age could suffer from the Simpson’s paradox^18^ due to the lack of overlap of age ranges across tissues, leading to false discoveries. To overcome these challenges, we developed a multi-level compositional analysis tool tailored to single-cell atlases by extending sccomp^19^ (Supplementary Methods) and incorporated multilevel differential expression analysis^12^ into tidybulk^20^.

Beyond these complexities and biases in data coverage, the presence of 19 cell-probing technologies and protocols represents a challenge in resolving biological effects (Supplementary results). While it is well known that single-cell technologies impact transcription quantification, it is unclear to what degree they affect cell abundance quantification. Principal component and differential analysis of cellular composition showed that while 10x protocols were consistent and did not show significant differences, Smart-seq2, microwell-seq^21^, and sci-seq did (Supplementary Figure S9). Estimating and accounting for the effect of technologies on tissue composition is therefore essential to test for biological differences and was part of our analysis pipeline.

### Ageing induces dysregulation of the immune system at the organ level

Age-related immune cell accumulation has been discussed broadly in the context of inflammaging^22^; however, it has yet to be systematically quantified across tissues and the human population. The harmonised data enabled us to track how immune cellularity (immune vs. immune) and composition change in ageing in different tissues. Using sccomp^19^, we estimated the effects of age on the immune system, accounting for other biological and technical effects, including sex, ethnicity, technology, study, and non-healthy status (see formula in Figure 4).

**Figure 4.**
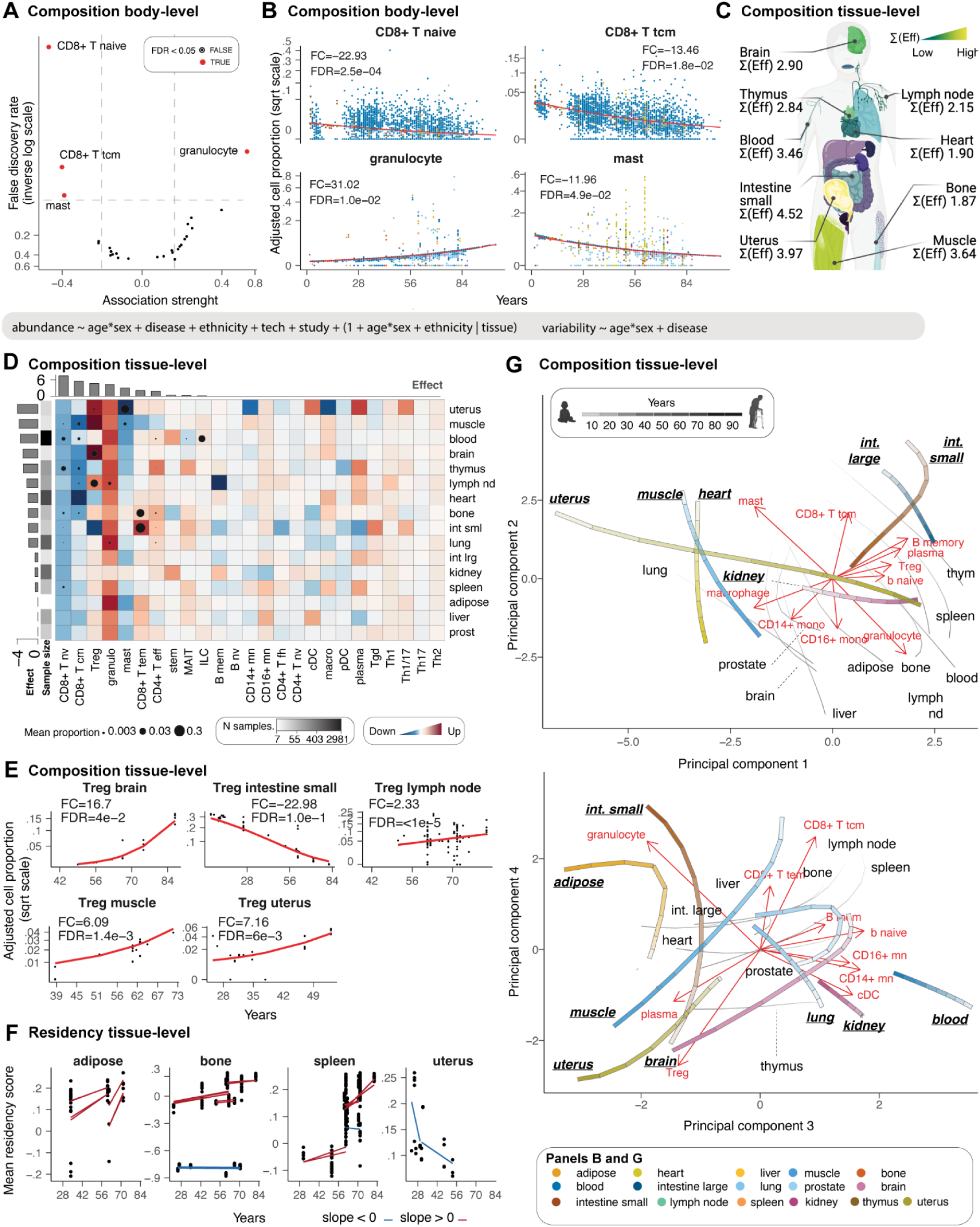
The landscape of immunity changes with age. A: Enriched and depleted cell types through age. **B:** Association between immune composition and age across tissues. Only cell types with significant associations (false-discovery rate < 0.05) are displayed. Points represent samples. The signed fold change represents the ratio between the estimated proportion of the age range limits present in the data (e.g. 39-73 for muscle). **C:** Average (across cell types) immune compositional changes associated with age for each tissue. “Eff” is the effect, represented by the coefficient of the slope. Yellow represents large compositional changes. Significant associations are labelled as text. **D:** Detailed view of age-related immune compositional changes for each cell type and tissue. The red colours of the heatmap represent a positive association. Cell types and tissues/tissue are colour-labelled as previously described. Histograms represent the magnitude of composition changes (sum of absolute significant effects) across tissue for rows and cell types for columns. **E:** Age-trends of Treg composition for several tissues, referring to Panel D. The signed fold change is explained for Panel B. **F:** Association between residency score of T cells and age. Red and blue represent positive and negative associations, respectively. Trend lines are shown across datasets. **G:** Principal Component Analysis, capturing the most significant variances in immune cell composition with age. The trajectories, represented by the colour-matched lines, indicate the direction and magnitude of changes in immune composition within each tissue over time, at ages 20, 40, 60, and 80 years, as denoted by the increasing intensity of the grey scale. The red text and arrows, termed ‘loadings’ in PCA, suggest the relative contribution of different immune cell types to the ageing trajectories within each tissue. The longer the arrow, the more that immune cell type contributes to the variance in the dataset along that principal component. We display the top ten.

We analysed individuals from 1 to 84 years of age. For immune cellularity analyses, we included 667 samples from 96 datasets and 17 harmonised tissues (Supplementary Table 1) that were not experimentally enriched or depleted for immune cells. For the immune compositional analyses, we included 6.296 samples from 189 datasets and 36 tissues (Supplementary Table 1). Each age group (10 years interval) and tissue included five samples on average for the cellularity analyses and 163 for the compositional analyses (Supplementary Table 2). The variability of the less represented factor combinations is learned from modelling the hierarchical structure of the whole dataset.

First, we sought to validate our observations on some well-known mechanisms. At the body level, we observed and quantified the well-documented^23–26^ decrease of naive CD8 T cells with age (Figure 4, panels A and B). We observed a reduction of naive CD8+ T cells of 22.93 folds (FDR <2.5×10^-4^) in an 8-to-84-year lifespan, validating known trends^27,28^. At the tissue level, we focused our validation on the thymus. While most immune compartments are dynamic and continuously change in response to infection or damage, the declining functional relevance of the thymus with age^29,30^ makes this primary lymphoid organ a suitable resource for validation. As expected, we identified the thymus as one of the most affected organs by ageing (Figure 4C). The ageing thymus was depleted in naive CD8+ T cells, with the largest effect across all tissues (Figure 4D) with a 4.19 fold change in a 2-61-year lifespan (FDR 5.0×10^-3^). On the contrary, CD4+ effector T cells were enriched with a fold change of 5.22 (FDR 1.2×10^-2^), representing the largest change among all tissues. These findings recapitulated and quantified well-known age-related phenotypic shifts of immune populations^25^ for the first time at the whole body scale.

Our analyses also identified unknown or less known ageing patterns. Despite inter-tissue heterogeneity, our model identified a significant body-level depletion of central memory CD8+ T cells of 13.46 fold change in an 8-to-84-year lifespan (FDR=1.8×10^-2^; Figure 2B)^31,32^. This trend was also significant at the tissue level (FDR<0.05; Figure 4D) for the thymus (FC=-10.43; FDR=5.6×10^-3^), blood (FC=-3.41; FDR=<5.0×10^-5^), lymph node (FC=-3.63; FDR=4.5×10^-3^), bone (FC=-10.65; FDR=1.1×10^-2^), heart (FC=-107.75; FDR=<5.0×10^-5^) and muscle (FC=-6.93; FDR=<5.0×10^-5^). Similarly, mast cells were significantly decreased at the body level with an 11.96 fold change in an 8-to-84-year lifespan (FDR=4.9×10^-2^), significant at the tissue level for the uterus and muscle, and noticeable in adipose tissue. On the contrary, granulocytes had a significant body-level increase with a 31.02-fold change in an 8-to-84-year lifespan (FDR 1.0×10^-2^; Figure 4B and 4D), which also shows tissue-level significance for lung, lymph node, and blood (FDR<0.05) and a large effect for adipose tissue. In contrast to the cell types showing conserved trends, T regulatory cells (Tregs) showed significant diversity in directionality across tissues (Figure 4D and 4E). For example, while they showed an age-related significant enrichment in the uterus (FC=8.12; FDR=6.1×10^-3^), brain (FC=17.64; FDR=4.0×10^-2^), lymph node (FC=2.35; FDR=<5.0×10^-5^), and muscle (FC=6.54; FDR=1.4×10^-3^), they were highly depleted in the small intestine (FC=-25.43; FDR=1.0×10^-1^) and, to a lesser extent, in the thymus and heart.

Motivated by the prevalence of T cells among the top compositionally dynamic cell types, we investigated the molecular contributors of T cell accumulation in tissues. We scored CD4+ and CD8+ T cells for residency^33^ (Figure 4F). Our results show that the increase in effector T cell infiltration in the small intestine and bone is linked to the up-regulation of a canonical residency signature^33,34^. We also observed this effect in adipose tissue, although the increase of CD8+ effector memory did not reach significance. On the contrary, the significant decrease of residency signature in the uterus is not mirrored by a T effector cell depletion.

In exploring the immune system’s ageing landscape, we used sccomp generative capabilities to chart the age-dependent trajectories of immune cell composition across a diverse array of human tissues. Utilising principal component analysis, we distilled the complexities of immune cell distribution into ageing programs (Figure 4G), revealing the nuanced interplay between universal and tissue-specific ageing patterns. Principal components 1 and 2 capture the primary two large-scale ageing programs (Figure 4G-top). The first is characterised by the compositional alteration of central memory CD8+ and mast cells, as opposed to granulocytes (mostly PC2), and the second by the enrichment of B cells and Tregs as opposed to macrophages/monocytes (mostly PC1). Most tissues, including the lymphatic and adipose, liver, prostate and brain, follow a shared main ageing program toward granulocytes and away from central memory CD8+ and mast cells. Among the tissues with an alternative ageing compositional program are small and large intestine. Notably, while the young small and large intestine share a similar immune composition, they diverge through ageing, with the mall intestine diverging away from Treg while approximating to macrophages/monocytes. The uterus is characterised by the largest transition, mostly along PC1 and mostly orthogonal to the small intestine, toward Tregs and the B cell compartment. The heart and skeletal muscle tissue show similar initial immune compositional, slightly diverging with age, with the muscle mostly aligning with the biggest tissue cluster, including lymphoid tissues. Principal components 3 and 4 (Figure 4G-top) describe a secondary ageing program driven by Tregs vs CD8+ T ratios, plasma vs. other B cells, and granulocyte vs. other myeloid. The largest cell-type cluster (grey lines in Figure 4G-bottom), primarily including lymphoid tissues, is characterised by a transition along the PC3, away from monocytes, classical dendritic cells, and B cells. The rest of the cell types are divided into two main trajectories. Muscle tissue, uterus and brain mostly follow the Treg-plasma direction, away from CD8+ T cells. In contrast, the small intestine mostly follows the granulocyte, CD8+ T cell direction and away from a monocyte/DC immune environment. Blood, lung and kidney, while dissimilar for the main ageing program in overall composition (location in the principal component space) and trajectory directions, show similarity in the secondary ageing program both in overall composition and trajectory. Notably, the trajectories of prostate and adipose tissue are similar, although their estimated compositional changes (Figure 4D) are minor.

These results provide a tissue-resolved reference of immune accumulation and composition through ageing. Our analyses highlight the power of our resource in systematically assessing key features of the immune system, such as cell-type-specific ageing patterns and the role of cell residency, as well as compositional ageing programs and trajectories that are conserved or tissue-specific.

### Ageing selectively influences immune cell communication and adaptability

Age-associated inflammation is often discussed as mediated by a low-grade persistent increase in inflammatory molecules, including soluble messengers^35,36^. To provide a quantitative landscape of age-related immune cell communication changes associated with significant compositional changes, we quantified immune cell ligand-receptor co-transcription within each sample^37^. Cell communication is quantified from single-cell RNA data based on similar transcription patterns between ligands and receptors across cell-type pairs^37^. Cell-communication analysis requires a relatively higher amount of information than compositional analysis, as a cell type must have sufficient cell and transcription abundance. Therefore, we focussed on tissues represented by a sufficiently large number of samples and with many cells per cell type, including blood, heart, lung, lymph node, kidney, and thymus. In total, we analysed 1,282 samples after filtering (see Methods). For the selected tissues, we quantified the average communication strength^37^ of each cell type versus all others, relying on well-known communication axes. We tested changes in communication strength through ageing using a custom Bayesian multilevel model^38^.

The relative loss of Tregs in the thymus (Figure 4D) co-occurred with the alteration of essential Treg communication patterns across lymphoid tissues (Figure 5A). Beyond communication axes that were down-regulated between Tregs and all other cell types, some communication axes, including those involving adhesion molecules, such as SELPLG and ICAM, were selectively downregulated between Treg and antigen-primed T cells (e.g. memory CD8+ T cells and CD4+ T cells), while they were upregulated in naive T cells (Figure 5B, Supplementary Figure S4).

**Figure 5.**
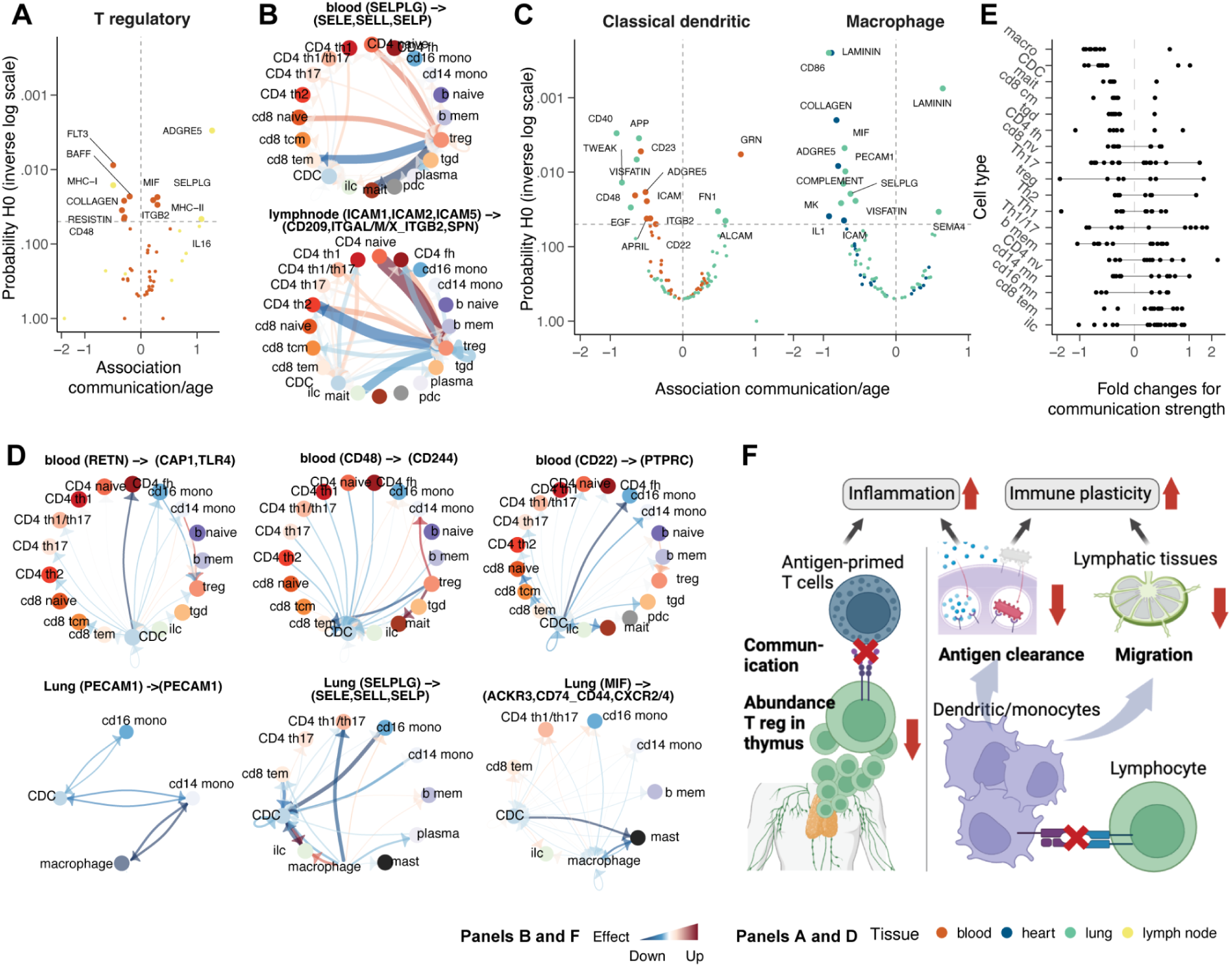
Changes in cell-cell communication with age. A: Association between communication axes (which underlie several ligand and receptor genes) transcription abundance and ageing for T regulatory cells. **B:** Estimated changes in communication strength across cell types through ageing, mainly in T regs. Red and blue represent increase and decrease, respectively. **C:** Association between communication axes transcription abundance and ageing for classical (myeloid) dendritic cells and macrophages. **D:** Estimated changes in communication strength across cell types through ageing, mainly in professional antigen-presenting cells. Red and blue represent increase and decrease, respectively. **E:** Overall communication changes across cell types. Dots represent communication axes. Lines represent the 95% credible interval. **F:** Cartoon of the major mechanisms of inflammaging and losing immune adaptability to new infections through ageing.

Macrophages and classical dendritic cells showed a consistent and strong decrease in strength across all significant communication axes (Figure 5C). Blood, heart and lung were the tissues for which we found the strongest evidence. The downregulated communication axes are mostly linked to dendritic cell migration and monocytic antigen clearance (e.g. ITGB2, ICAM, PECAM1, SELPLG and MIF), antigen presentation^39^ (e.g. RETN), cytokine production in dendritic cells^40^ (e.g. CD48). The downregulated monocyte communication (Figure 5D, Supplementary Figure S4) primarily targeted T cells (e.g. for RETN, CD48) and other monocytes (e.g., for CD22, PECAM1, SELPLG, and MIF).

Given these findings, we sought evidence of an overall depletion of immune cell communication in ageing. Analysing all significantly changed communication axes across all cell types (Figure 5E), we observed an almost exclusive depletion for several cell types, including macrophages, classical dendritic cells, MAIT cells, CD8+ central memory, CD4 helper and gamma-delta T cells. In contrast, we observed no cell type with the opposite trend. These changes pointed toward an immune-system-wide age-related loss of immune cell communication.

Our results highlight the age-related impaired cell communication of Tregs and myeloid cells in multiple tissues, likely contributing to loss of immune specificity and adaptability to new infections (Figure 5F). The emerging gene programs are related to the regulation of inflammation, decreased migration, antigen clearance, and antigen presentation.

### Sexual dimorphism shapes the immune system in a cell-type-specific manner

Sexual dimorphism is a well-known phenomenon related to differential susceptibility to disease and response to treatments^3^. To characterise the sex-related differences in the immune system across tissues at the cellular and molecular levels, we performed multilevel analyses contrasting female and male populations while accounting for other biological and technical effects (i.e. age, ethnicity, technology, study, and non-healthy status; see formula in Figure 6). We analysed individuals with available sex information and filtered for tissue represented in both sexes from individuals older than eight years old. For cellularity analyses, we chose samples with no experimental immune enrichment or depletion. We filtered the data for high quality and replication (see Methods), obtaining 469 samples from 89 datasets and 10 harmonised tissues (Supplementary Table 1). For composition analysis, we used 4,588 samples from 63 datasets and 14 tissues (Supplementary Table 1), for which we only selected the immune cells. On average, each sex group and tissue included 23 samples for the cellularity and 163 for the compositional analysis (Supplementary Table 2).

**Figure 6.**
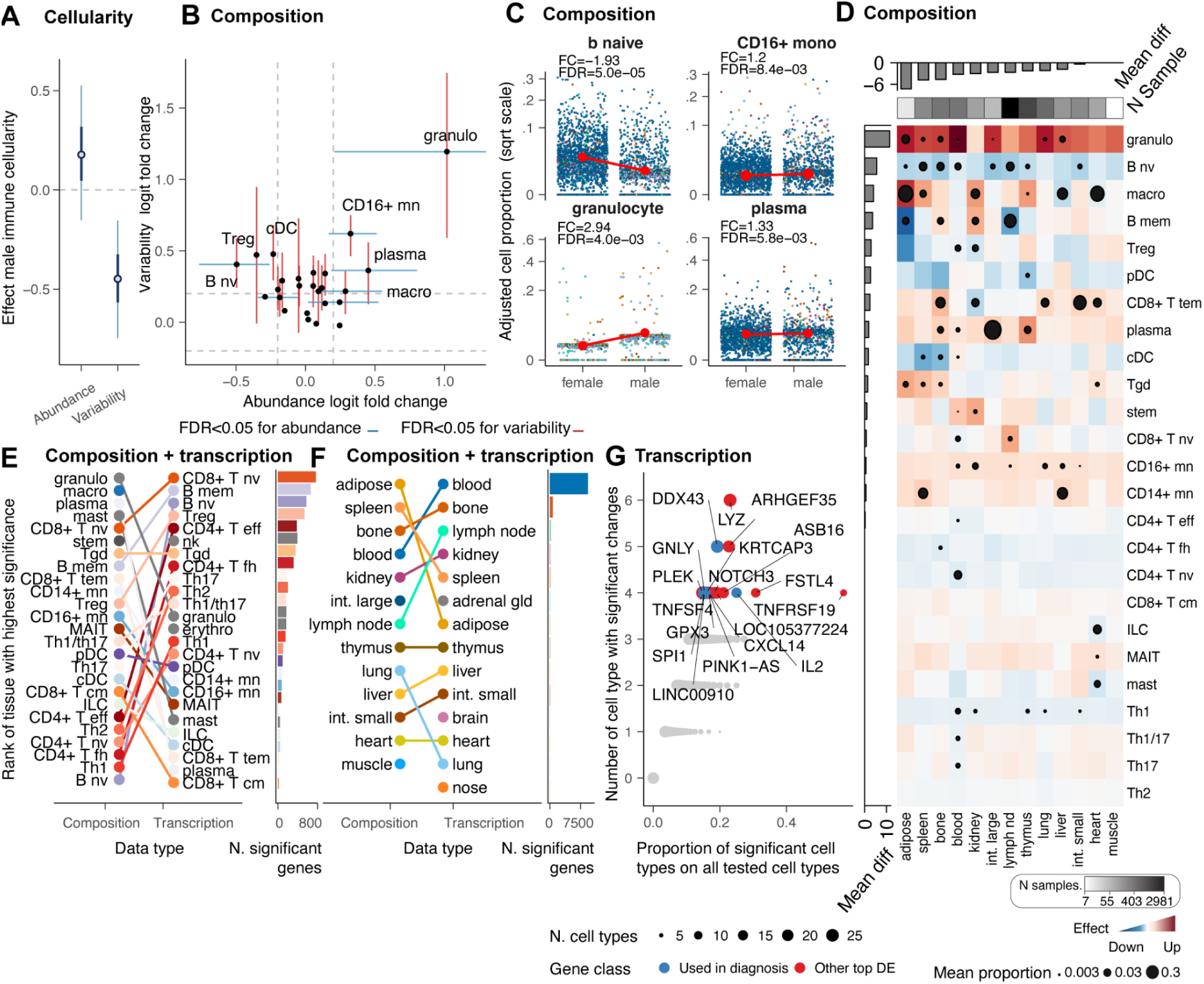
Sex-related changes in immune cell composition and transcription. A: Effect of sex on immune cellularity and its variability. The thin and thick lines represent the estimated effect’s 95% and 60% confidence intervals, respectively. The dot represents the point estimate. **B:** Compositional (blue) and variability (red) differences at the body level for males compared with a female baseline. Points represent mean effects, and error bars represent the 0.95 credible intervals for significant associations. **C:** Association between immune composition and sex across tissues. Only cell types with significant associations (false-discovery rate < 0.05) are displayed. Points represent samples. The signed fold change represents the ratio between the estimated proportion of the age range limits present in the data. **D:** Immune composition differences per tissue. Red and blue represent enrichment and depletion in males compared with a female baseline. Dots represent significant associations (FDR < 0.05). The dot size represents significance probability; the bigger, the higher, the lower the FDR. The signed fold change represents the ratio between the estimated proportion of the age range limits in the data. **E:** Cell-type rank for the overall highest significance (across tissues) for compositional and transcriptional differences. **F:** Tissue rank for the overall highest significance (across cell types) for compositional and transcriptional differences. **G:** Most altered genes across tissues and cell types (estimated with a multilevel model for each tissue, with cell types as a grouping variable). Black circles are displayed only for significant associations (FDR<0.05), and their size depends on the cell type abundance.

We estimated no significant body-level difference in mean immune cellularity between sexes, with the female cohort appearing to be more variable than the male (Figure 6A). However, several compositional differences emerged at the body level (Figure 6B and 6C), including B naive, plasma, monocytes (CD16+), and granulocytes (FDR<0.05). The deconstruction of compositional differences across tissues validated known aspects of immune sex dimorphism. These include the enrichment of the CD4+ T population in females^41–43^, which we observe for CD4+ naive, helper 1, follicular helper and effector, and a male enrichment of CD8+ effector memory^41–43^. Our compositional body map also shows that most locations were dimorphic (Figure 6D). The adipose tissue had the highest effect (mean across cell types), followed by the spleen and bone marrow. While the trends of granulocytes, B cell naive, plasma and CD16+ monocytes are consistent across tissues, reflecting the body-level findings (Figure 6C), Tregs, memory B cells and macrophages are highly heterogeneous across tissues with opposite trends depending on the body location. For example, while macrophages are significantly enriched in males in the adipose tissue, they are significantly enriched in females in the heart and liver (Figure 6C).

To investigate potential phenotypic differences associated with compositional changes, we performed a differential gene-transcript abundance analysis among 116,606 cell-type-resolved pseudobulk samples (see Methods). We tested differences in transcriptional abundance between sexes, taking into account other biological and technical effects^12,20^. Overall, the compositional differences appear orthogonal to the transcriptional differences (Figure 6E). While myeloid cells tend to be distinct between sexes at the compositional level, the lymphoid compartment has the largest transcriptional differences. Among the lymphocytes with the most transcriptional changes are naive CD8, T regulatory, CD4 effector, follicular helper and gamma delta. The dichotomy between compositional and transcriptional changes was less present in tissue-level analyses (Figure 6F), where lymphoid tissues, including bone, spleen, and lymph nodes, are some of the most compositionally and transcriptionally distinct.

Despite this heterogeneity, we sought to identify the significance and clinical impact of global transcriptional changes conserved at the tissue and cell-type levels (Figure 6G). Genes that were most consistently differentially transcribed included several immunotherapy targets. IL2 is targeted in immunotherapy, particularly in the treatment of certain cancers like melanoma and renal cell carcinoma^44^; the immune genes TNFSF4, important in T cell activation^45^, have been proposed as immunotherapy targets, and GNLY, important in NK cell biology, is a diagnostic marker and have potential as immunotherapy target^46^

Our results resolve the tissue and cell-type specificity of sex-related immune differences, ranking naive B cells and granulocytes as most consistently affected across tissue and immunotherapy gene targets among the most affected overall.

### Sex-differential ageing has a dominant role in immune sexual dimorphism

To understand whether the effects of ageing and sex on the immune system are intertwined, we tested the significance of sex-dependent differential ageing at the cellularity, composition, and transcriptional levels. Differential ageing refers to age-related changes in the immune system, the rate of which differs between females and males. For example, a gene transcript or cell type is enriched in ageing for females but depleted for males.

We performed transcriptional analyses on 116,606 cell-type-resolved pseudobulk samples using our model, which captures the age-sex interaction^12,20^ (see equation in Figure 4; see Methods), also accounting for the effect of sex and age independently from each other and other biological and technical factors. First, we compared the number of genes for which transcription was associated with differential ageing or with sex only (strictly independently of age). We observed that sex dimorphism was dominated by differential ageing (Figure 7A), with 294 significant genes (across more than five tissues) versus 246 genes associated with sex only. The gene overlap between the two effects was minimal (23 significant genes). This finding motivated further compositional and transcriptional analyses aimed at resolving cellular and transcriptional drivers.

**Figure 7.**
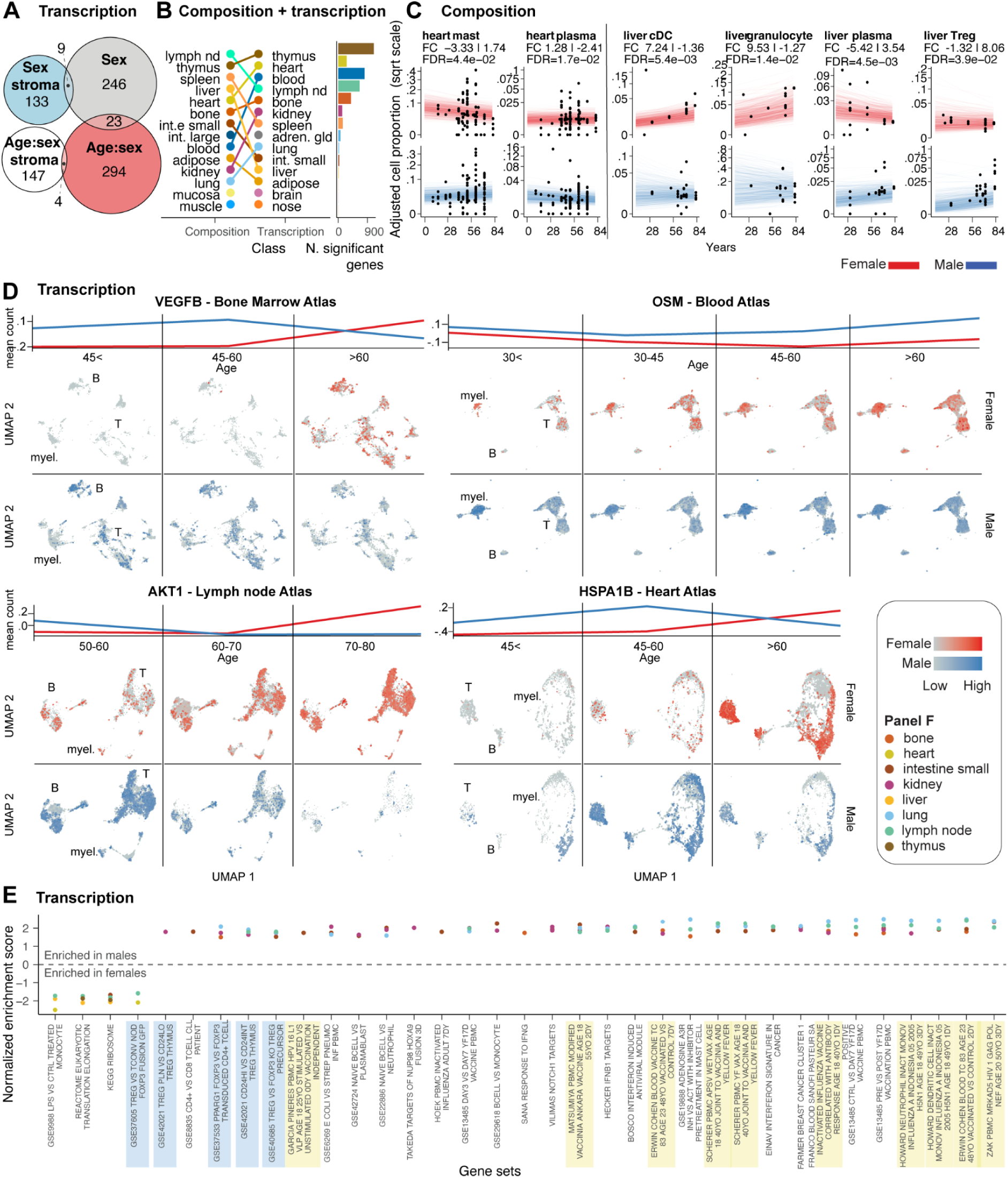
Sex-related differential ageing of the immune system. A: Overlap between genes in immune cells whose transcription abundance is significantly associated with sex or differentially associated with age depending on sex (“age:sex”; increasing during ageing in males while decreasing in females). Stroma indicates non-immune cells. **B:** Tissue rank for the overall highest significance (across all cell types) for compositional and transcriptional differences. **C:** Differential ageing for the immune composition of two non-lymphoid tissues more impacted by differential ageing. For each tissue, cell types are shown for which females and males showed significant differences with opposite trends (e.g. increasing through age for females but decreasing for males). Each dot represents a sample. Each line represents all possible predicted trends from the model, indicating the level of uncertainty. The signed fold change represents the ratio between the estimated proportion of the age range limits present in the data. The first displayed fold change is for females, while the second is for males. **D:** Age-resolved UMAPs represent significantly associated genes for the most altered tissues. The top represents the differential trends between sexes, along the binned age axis, expressed in mean (across samples) normalised count. **E:** Gene set enrichment analysis. The top 30 gene sets (for average normalised enrichment score), altered in at least three tissues (colour-coded), are shown. Blue-shaded sets include immune regulation. Yellow-shaded sets include age-dependent responses to vaccination.

Immune cellularity at the body or tissue level (analysed across 469 samples after filtering) and its variability did not appear to be strongly associated with ageing depending on sex (Supplementary Figure 3). Similarly, immune composition at the body level showed no significant association (Supplementary Table 3). However, we identified significant differential ageing at the tissue level (Figure 7B; Supplementary Figure 3), most notably in the lymphoid tissues and the heart and liver. Generally, we observed that the differential ageing for specific cell types is heterogeneous across tissues. For example, focusing on the heart and liver (Figure 7C), we estimated plasma cells becoming relatively more abundant in ageing females in the heart but in males in the liver. In the heart, we observed another large effect in mast cells being relatively enriched in ageing males (FC difference of 5.07, FDR=4.4×10^-2^). In the liver, Tregs were relatively enriched in ageing males (FC difference of 9.38, FDR=3.9×10^-2^), and cDC (FC difference of 8.60, FDR=5.4×10^-3^) and granulocytes (FC difference of 10.80, FDR=1.4×10^-2^) in ageing females.

We then focused on the tissues highly affected by differential ageing both transcriptionally and compositionally for more in-depth analyses at the single-cell level. We composed single-cell atlases of the bone marrow, the blood, the lymph node and the heart to resolve the cellular heterogeneity of the top affected genes (Figure 7D). As a common trend among these tissues, we observed sex-dependent differential age trajectories of key inflammatory and cancer-related genes. For example, VEGFB, AKT1, and HSPA1B in the bone marrow, lymph nodes, and heart atlases were upregulated in men during middle age (up to 60 years) and downregulated later in life. On the contrary, gene upregulation can be observed only in older females (up to 80 years). A different pattern is shown by the gene OSM1, which may be involved in cellular processes related to oxidative stress and inflammatory responses in cancer^47,48^. OSM1 shows a consistently higher gene transcript abundance in males across all age groups examined and with a differential age trend compared to females.

To gain a higher-level perspective on gene programs related to differential ageing, we performed a gene set enrichment analysis across tissues, using the rank test in GSEA through tidybulk^20,49^. Among the 30 top significant pathways altered in at least three tissues, signatures related to age-related differential response to vaccination were recurrent, including yellow fever, human papillomavirus, and influenza (Figure 7E). Similarly, recurrently altered pathways included immune regulation.

Our results show that sex-dependent differential ageing of the immune system was a significant component of sex dimorphism. We rank the heart and kidney as the most affected organs for immune cellularity and several lymphoid tissues as significantly transcriptionally altered for several cancer-related genes.

### Ethnic group heterogeneity affects diagnostic and therapeutic immune targets

The immune system evolved to protect us from environment-specific pathogens^50^. These forces create transient and genetic immunological differences among ethnic groups^51^. To resolve ethnic-specific immune characteristics across the human body at the cellular and molecular levels, we compared the major ethnic groups for immune cellularity, composition and transcriptional programs.

For cellularity analyses, we chose samples with no experimental immune enrichment or depletion. We filtered the data for high quality and replication (see Methods), obtaining 667 samples from 96 datasets and 17 harmonised tissues (Supplementary Table 1). For composition analysis, we used 6,293 samples from 189 datasets and 36 tissues (Supplementary Table 1), for which we only selected the immune cells. On average, each sex group and tissue included 23 samples for the cellularity and 65 for the compositional analysis (Supplementary Table 2). A total of 5,880 samples (in total across analyses) were derived from Europeans, 790 from Asian descendants, 273 from Africans, 115 from Hispanic ethnicity, and 3,852 by unknown origin (Figure 8F). As the European ethnicity was the most represented, we selected it as a baseline and sought cellular and molecular immune differences against other ethnicities.

**Figure 8.**
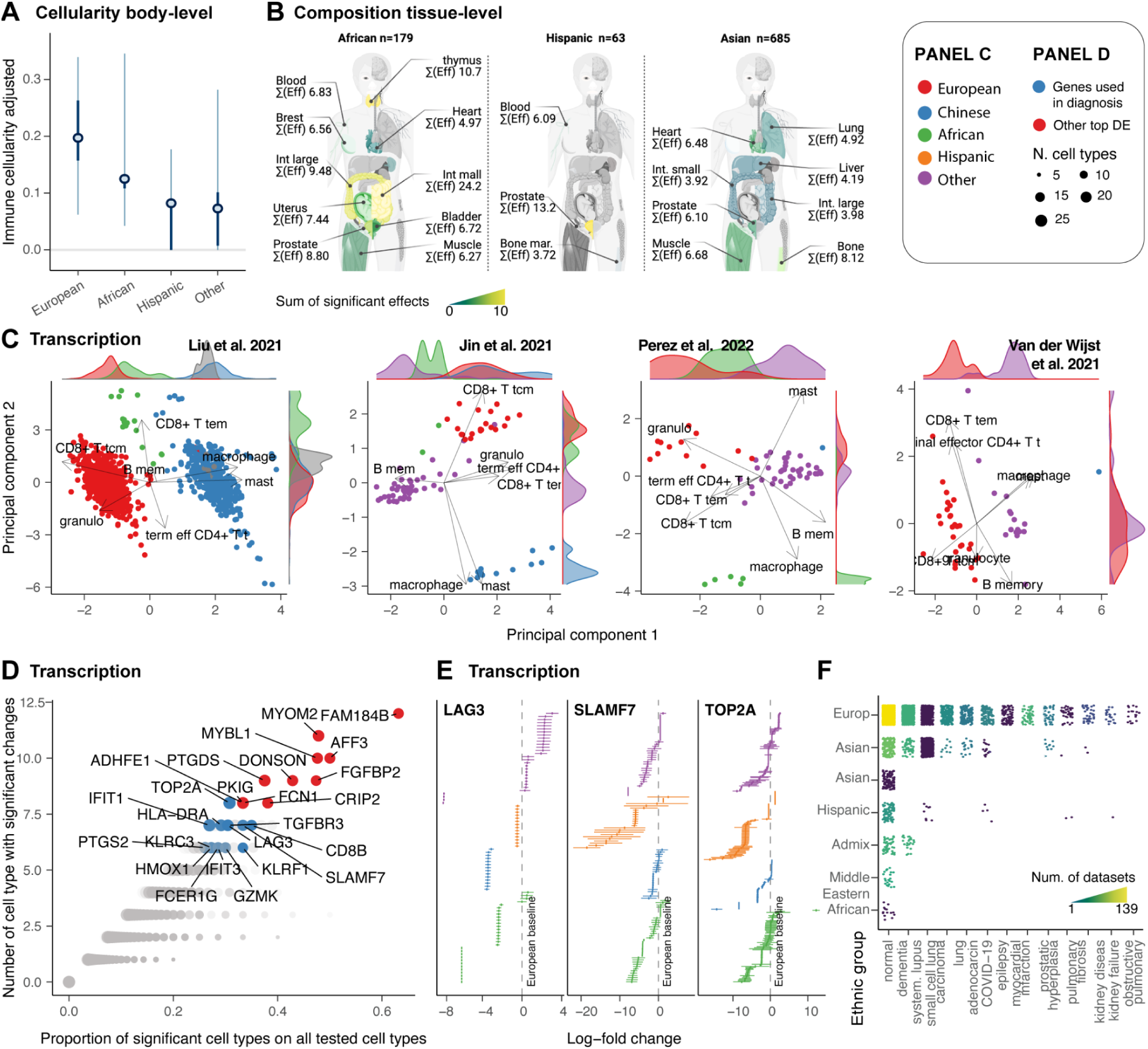
Changes in immune cell composition with sex and ethnicity. A: Overall effect of sex on immune cellularity and its variability with females as the baseline. The mean, 50% and 95% credible intervals are visualised. **B:** Tissue-specific differences of immune cellularity between sexes. Left: cartoon of the human body coloured by significant effect. Right: Immune cellularity of samples, with non-sex effects removed. **C:** Sample distribution along reduced dimensions (principal components) coloured by ethnicity. **D:** Gene transcripts differentially abundant between Europeans (baseline because highly represented) and other ethnicities across several tissues and cell types. **E:** Credible intervals of selected top targets among the differentially abundant gene transcripts. **F:** Representation of the sample size for ethnicity across diseases. Dots represent samples. Quadrants are coloured according to the number of datasets.

For overall immune cellularity, Europeans showed a higher mean value (21%, non-significant) compared with Africans and Hispanics (<15%, Figure 8A). Compositionally, at the tissue level, we detected several significant differences (Figure 8B). Ranking the ethnicity-tissue pairs with the highest mean effect (across cell types, only considering significant changes) put the African ethnicity as the most distinct from Europeans, followed by Asians. Hispanics, as expected, were the less distinct. For the African ethnic group, the intestine, thymus and the prostate were the most impacted tissues. The prostate was ranked among the top impacted tissue for all three ethnicities.

To characterise the heterogeneity of immune composition across ethnicities, we mapped the adjusted proportions of immune cells (excluding non-ethnicity variation) onto principal components. We focused on blood, the most data-rich tissue, and the largest datasets with multiple ethnicities^52–55^. Across all datasets, we observed marked clustering of samples from the same ethnicity along principal components 1 or 2 (Figure 8C). The cell types contributing most to that variability are memory CD8+ T cells and granulocytes, characterising the Europen population, and macrophages and mast cells, characterising the Chinese population.

Then, to understand the potential clinical impact of this diversity, we tested transcriptomics differences across ethnicities and sought the presence of therapeutic targets or diagnostic markers among the most impacted genes. We focused on the most frequently altered genes across cell types (and across five or more tissues; Figure 8D). A core of 23 genes was significant in 30% to 60% of the cell types tested against (Figure 8D). Of those, 13 are or have been proposed as diagnostic markers or immunotherapy targets (TOP2A^56^, LAG3^57^, SLAMF7^58^, HLA-DRA^59^, IFIT1 and IFIT3^60,61^, PTGS2, HMOX1^62^, GZMK^63^, FCER1G, KLRC3 and KLRF1^64^, TGFBR3). Of particular relevance (Figure 8E), Topoisomerase II alpha (TOP2A) is a target for anti-cancer drugs^56^, LAG3 is a checkpoint inhibitor^57^, SLAMF7 is targeted by the myeloma immunotherapy drug Elotuzumab^65^, PTGS2 is a target for anti-inflammatory drugs with a role in inflammatory diseases and cancer^66^.

Our results characterise and quantify a strong effect of ethnicity on the immune system at the cellular and molecular levels. This heterogeneity includes key diagnostic and treatment gene targets, which are especially relevant considering the under-representation of non-European ancestry.

## Discussion

This study comprehensively mapped the immune system across tissues and the human population. Compared with previous multi-organ studies^6–8^, we significantly increased tissue and human population coverage, which allowed us to untangle demographic effects and interactions at the body and tissue levels using sccomp. Modelling immune composition, transcriptional activity and cell communication, we uncovered conserved and tissue-specific immune-ageing programs, significant changes across sex, how sex determines specific immune-ageing profiles, and significant heterogeneity across ethnicities. We share these data with the community through a programmatic query interface in R and Python, *CuratedAtlasQuery*.

To deal with the complex structure typical of atlas-level datasets, we developed a method for multilevel compositional analysis. This method extends sccomp^19^, a robust and flexible Bayesian model able to estimate differences in relative abundance and variability in an outlier-insensitive manner. Our method, applied to this large-scale dataset, allowed the extraction of age, sex and ethnicity-related effects nested within tissue and study effects. It prevents biased estimation favouring over-represented groups (i.e., tissues and studies) at the expense of under-represented ones. This tool can model complex effects at tissue and body levels, representing a scalable tool for atlas-level analyses.

In our healthy immune cell map, age emerged as a strong predictor of composition. The ageing of the immune system, especially for lymphoid tissue, has been widely discussed in the literature^36^. This study extends the current knowledge by contributing a compendium resource with a holistic quantification of the ageing effects in the context of non-lymphoid tissues, highlighting what aspects are conserved at the body level or tissue-specific. Notably, the shift from naive to antigen-experienced lymphocytes is a finding that aligns with and validates previous research^25,67^. This loss represents one of the tissues’ most significant global effects. Lymphoid tissues were among the most altered with age, including the thymus, recapitulating the well-known progressive atrophy^29,30^ and providing further evidence for the robustness of our study. Our findings suggest tissue-specific T-cell accumulation may not always correlate with residency pathways, posing questions for future research, especially regarding the uterus’s unique immune alterations with age. For example, for the uterus, whether a continuous influx and efflux drives the immune cell accumulation in these tissues requires future investigation. Among all tissues, the uterus is the most affected by age in terms of immune composition. The relationship between ageing and the immune composition of the uterus is complicated by the effects of menopause, chronic viral infections, microbiota and lifestyle factors^68,69^. Nonetheless, quantifying the homeostatic trajectory for immune composition can serve as a calibration tool for future studies.

Although we identified several cell types that share changes across tissues, we observe a range of cell types for which ageing trends are tissue-specific. Among those cell types, Tregs had a strong compositional association with age but with diverse directionality based on the tissue. This striking pattern indicates that Treg compensation of age-related inflammation might be a local, tissue-specific process rather than a homogeneous phenomenon^70,71^. Characterising this heterogeneity further can inform immunotherapy for cancer and organ transplants. We observed differential Treg ageing pathways between tissues, which are also across the cell-communication pathways (Figure 5A). Although the list of tissues with enough data availability to allow communication analyses is limited, the dichotomy we identified supports our findings for Tregs at the compositional level. Overall, altered cell composition and communication patterns indicate a major and multifaceted role of Treg suppressive function, migration, and localisation in the ageing immune system^72^.

Our model allowed us to quantitatively track tissue-specific ageing trajectories and discover body-level and tissue-specific immune ageing programs (Figure 4G). This temporal map resonates with previous efforts to produce immune ageing metrics^73^. However, our analyses go beyond the concept of cell-type specific immune ageing, resolving tissue-specific ageing programs. We showed that while a larger tissue group includes the lymphoid and several non-lymphoid tissues, such as lung, adipose and liver, which follow a similar ageing trajectory, tissues such as the uterus, kidney and small intestine have orthogonal evolution.

This orthogonality is also present in the large and small intestine, suggesting that immune ageing heterogeneity exists between associated tissues. This map contrasts with the idea that tissues converge during ageing toward a loss of tissue identity^74^. We showed further complexity with a secondary ageing program, characterised by the balance of cell states within each major cell type (e.g., plasma vs non-terminally differentiated B cells). In this program, blood and other highly vascularised tissues diverge from other lymphoid tissue. These trajectories could serve as an age reference to provide an immunological clock of tissues in healthy and diseased states.

A substantial depletion of cell communication also emerged for professional phagocytes, including macrophages and dendritic cells in the lungs and blood (Figure 5F). The disrupted communication axes included antigen presentation and tissue trafficking. This observation supports the idea that decreased antigen clearance by macrophages and dendritic cells is a critical mechanism in ageing^75–77^. Concurrent with the down-regulation of pathways linked to the ability of myeloid cells to clear and present antigens to adaptive immune cells, we observed a significant and global increase in monocyte infiltration across tissues (Figure 4 E and F). We provide clarity over the controversy of whether aged dendritic cell subtypes demonstrate specific alterations in antigen presentation capacity^25^. Age-related inflammation (inflamaging^78^) is a well-studied phenomenon^25^; however, its investigation is spread across clinical areas without a systematic and quantitative investigation at the population and tissue levels. Our age-resolved tissue map of the immune system supports the idea that inflammaging and loss of adaptability to new infections are intertwined and synergic. As non-linear transcriptional patterns have been identified in ageing mice^67^, future work will develop flexible tools able to fit non-linear effects to multilevel, atlas-level compositional and transcriptional analyses.

Sex dimorphism has been characterised at the molecular level across diseases. For example, X-linked genes and sex hormones play significant roles in autoimmune diseases^3^. Cancer incidence, progression and therapy response differ between sexes due to dimorphism at the hormone levels and gene expression^79^. Sex-related genetic differences influence drug metabolism, distribution, and excretion^80^. Immunologically, hormones such as estrogen can up-regulate immune responses, while testosterone may have immuno-suppressive effects^3^. Testing for sex-related differences in immune cellularity, composition and transcription across tissues, independent of age and ethnicity, we identified differences in the lymphoid and non-lymphoid compartments, aligning with public knowledge^3^. Multiple studies^41–43^ identified the CD4+ T population enriched in females independently of ethnicity. While this finding matches our results, our analyses further resolved the subtypes with homogeneous trends across tissues, including T helper 1 (Th1), follicular helper, and naive CD4+ T cells. Our result translates the evidence for a female enrichment of Th1 in infection models^81^ to human data. We observed a slight male enrichment of Th17 and 1/17 for most tissues. Similarly, consistent with the literature^41–43^, we observed a male enrichment of CD8+ effector memory T cells in most tissues except for the kidney. The predominance of B-cell transcriptional changes previously observed in blood^82^ is observed in our data across all tissues (Figure 5E).

Our results highlight the heterogeneity of Treg sex-proportionality differences across human tissues, supporting a multifaceted role of this cell type between sexes. The complexity of the role of Tregs in tissue biology is reflected in the literature, presenting contrasting evidence about Tregs’ proportionality of females vs males between human and model organisms^83^. For example, analysing 3,210 blood samples, we provide contrasting evidence to previous observations^84^, supporting a significant enrichment of Treg in the blood of females. Importantly, we show that sexual dimorphism can have practical clinical implications, with some of the most differential genes across tissues and cell types being established or proposed immunotherapy targets, such as IL2^44^; TNFSF4^45^, and diagnostic markers, such as GNLY^46^.

Our results highlighted that a significant contributor to immune sexual dimorphism is differential ageing, with an equivalent number of affected genes across tissue and cell types. In contrast with ageing-only and sex-only effects for which global body-level changes exist, differential ageing appears to be primarily tissue-specific. Compositionally, besides lymphoid tissues, the heart and the liver are the most affected tissues. The single-cell analyses of tissue atlases, with the largest overall transcriptional differences, reveal distinct gene regulation patterns across tissues. These genes are markedly connected to tumorigenesis^85^, tumour growth^86^, cancer progression^87^ and metastasis^88,89^. These findings strengthen the idea of the need for personalised cancer treatment strategies based on the age and sex of the patients. Differences in vaccination immune response emerged as highly enriched in our top differential gene sets. Those signatures, originally linked to age differences in vaccination, include many genes we more specifically link to differential ageing.

Our analyses draw a prominent picture of data production bias toward European ethnicity. This bias might have important implications for precision medicine research in case clinically relevant features of the immune system showed marked heterogeneity across ethnic groups. Our analyses support a well-known immune diversity across ethnicities^4,5^. Our large-scale analysis adds tissue resolution to an established connection between ethnic and geographic diversity and the immune system^51^. A stark diversity emerged compositionally at the tissue level, with the African ethnicity showing the largest differences from Europeans compared to other ethnic groups. These differences were concentrated in the thymus, intestine, prostate, and breast. Transcriptionally, we produced several lines of evidence for diversity across ethnicities (Figure 8). Notably, many top genes consistently differentially expressed across tissues and cell types for non-European ethnicities are clinically important. For example, TOP2A, LAG3, SLAMF7, and PTGS2 are established or promising key targets for cancer treatments and immunotherapy^56,57,65,66^. Independently of these differences being linked to genetics or geographical origin, this finding represents a call for a more inclusive and balanced data collection strategy across ethnic groups, enhancing precision medicine in the coming years^90^.

Our study employed meticulous design and advanced analytical tools to minimise the influence of outliers and ensure robustness against immune fraction enrichment or depletion. Nonetheless, caution must be exercised when interpreting results from specific tissues. For instance, estimating immune cellularity in non-lymphoid tissues faces challenges due to the inevitable depletion of stromal and other cell types during tissue processing and library preparation. Additionally, blood contamination in solid tissues like the lungs and heart introduces a layer of variation that warrants consideration. The inherent difficulties in sampling specific immune cell types, such as granulocytes, with single-cell technologies are also acknowledged. However, these factors do not impact the differential analyses, as our statistical model includes those factors and accounts for tissue and study cell-specific extraction biases. Recognising the constraints in drawing inferences for underrepresented groups^91^, our interpretations of highly resolved patterns, such as tissue-specific compositional changes across different demographics, are deliberately conservative. For example, we avoided interpreting the effects of specific cell types/tissue combinations for specific underrepresented ethnic groups and instead focused on overall heterogeneity or transcriptional trends common to many tissues and ethnic groups.

Our map of immune cellularity, composition, transcription and cell communication captures the intricate immunological landscape across the diverse human population. The heterogeneity at the human population level is rarely incorporated into the study of markers of disease and therapeutic targets. Using a Bayesian model to account for the complex dependence structure of the data, we resolved changes in the immune system associated with age, gender, ethnicity and technology. A key contribution of this work is to make single-cell data in the Human Cell Atlas more accessible to users in a curated form via our CuratedAtlasQuery interface and provide a quantitative immune system model. This resource has the potential to provide groundbreaking insights into the healthy immune system and function as a baseline for detecting disease-driven loss of homeostasis. By sharing our immune map with the scientific community, we hope to catalyse further breakthroughs in cancer, infectious disease, immunology and precision medicine.

## Data availability

The count data used for this study can be downloaded using the query interface *CuratedAtlasQuery*, which can be found in Bioconductor and at github.com/stemangiola/CuratedAtlasQueryR. The proportion estimates can be found at github.com/stemangiola/immuneHealthyBodyMap

## Code availability

The code used for the analyses and the figures can be found at github.com/stemangiola/immuneHealthyBodyMap. Our multilevel model can be found at github.com/stemangiola/sccomp/tree/multilevel.

## Acknowledgements

We thank the Human Cell Atlas Consortium for the CELLxGENE data source. We thank the Chan Zuckerberg Initiative for formatting and organising the data used in this study. We thank all individuals who donated tissues. We acknowledge all WEHI’s ITS and Research Computing Platform members for their computing support. We thank the Stan community for their constant support. This research was supported by the Nectar Research Cloud, a collaborative Australian research platform supported by the NCRIS-funded Australian Research Data Commons (ARDC). The Novo Nordisk Foundation Center for Stem Cell Medicine, reNEW, is supported by a Novo Nordisk Foundation grant number NNF21CC0073729.

S.M. was supported by the Victorian Cancer Agency Early Career Research Fellowship (ECRF21036). A.T.P. was supported by an Australian National Health and Medical Research Council (NHMRC) Investigator Grant (2026643). S.M. and A.T.P. were supported by the Lorenzo and Pamela Galli Next Generation Cancer Discoveries Initiative. A.K. was supported by an NHMRC Investigator grant (2017420). S.B. was supported by the National Health and Medical Research Council of Australia (2008408), a 350th Anniversary Research Grant from Merck KgGA, The Advanced Genomic Collaboration, and the International Research Training Group (IRTG2168) funded by the German Research Council and The University of Melbourne. M.M. was supported by the NHGRI and NCI of the National Institutes of Health under award numbers U41HG004059 and U24CA180996, and by the Chan Zuckerberg Initiative DAF, an advised fund of Silicon Valley Community Foundation. F.J.R. receives institutional support as a coinvestigator and is subcontracted by the Peter MacCallum Cancer Centre for an investigator-initiated trial, which receives funding support from Sanofi/Regeneron Pharmaceuticals. V.J.C. was supported by the grant “Bioconductor: An Open-Source, Open-Development Computing Resource for Genomics” NIH NHGRI Project Number2U24HG004059-17. The research benefited from the Victorian State Government Operational Infrastructure Support and Australian Government NHMRC Independent Research Institute Infrastructure Support.

## Contributions

SM conceived, designed and implemented the study and performed the analyses under the equal supervision of ATP, Martin M, SB and AK; Michael M. and YE implemented the CuratedAtlasQuery interface; NR, CL and WS performed data analyses for Figures 1 and 2. VY and BP contributed to revising the analyses and the manuscript. WH supported data curation. All authors contributed to the article’s writing.

## Star Methods

| REAGENT or RESOURCE | SOURCE | IDENTIFIER Software and algorithms |
| --- | --- | --- |
| Software and algorithms |  |  |
| sccomp | Mangiola et al.,<sup>19</sup> | https://github.com/stemangiola/sccomp/commit/1ceefdbbf6873f09af7811f6791b3f9e4aefb163 |
| CuratedAtlasQuery |  | https://github.com/stemangiola/CuratedAtlasQueryR/commit/338c973250332cdbde80ee6ca56cc6c7951152d1 |
| tidybulk | Mangiola et al.,<sup>20</sup> | https://github.com/stemangiola/tidybulk/commit/58d34393d0dd970f663d0ba31b0248c0fad50462 |
| glmmSeq | Lewis et al.,<sup>12</sup> | https://github.com/myles-lewis/glmmSeq/commit/8a473e1565a975e34263151ad9265dc6f469dae8 |
| brms | Burkner,<sup>13</sup> | https://github.com/paul-buerkner/brms/commit/701a6993f8a5e8b65799ee3b333e05d31a10219f |
| cellxgenedp |  | https://bioconductor.org/packages/release/bioc/html/cellxgenedp.html#:~:text=cellxgenedp_1.6.1.tar.gz |
| Azimuth | Hao et al.,<sup>92</sup> | https://github.com/satijalab/azimuth/commit/243ee5db80fcbffa3452c944254a325a3da2ef9e |
| SingleR | Aran et al.,<sup>16</sup> | https://bioconductor.org/packages/release/bioc/src/contrib/SingleR_2.4.1.tar.gz |
| celldex |  | https://bioconductor.org/packages/release/data/experiment/src/contrib/celldex_1.10.0.tar.gz |
| CellChat | Jin et al.,<sup>37</sup> | https://github.com/sqjin/CellChat/commit/e4f68625b074247d619c2e488d33970cc531e17c |
| singScore | Bhuva et al.,<sup>93</sup> | https://github.com/DavisLaboratory/singscore/commit/903177c62e42fff001fe4fdf0b34fc5d3bf924a5 |
| edgeR | McCarthy et al.,<sup>94</sup> | https://bioconductor.org/packages/release/bioc/src/contrib/edgeR_4.0.1.tar.gz |
| Seurat | Butler et al.,<sup>95</sup> | https://github.com/satijalab/seurat/releases/tag/v5.0.1 |
| tidySummarizedExperiment | Hutchison and Keyes et al.,<sup>96</sup> | https://github.com/stemangiola/tidySummarizedExperiment/commit/f249098b4bbf1a7e57c2e211555ee277421c5553 |
| tidySingleCellExperiment | Hutchison and Keyes et al.,<sup>96</sup> | https://github.com/stemangiola/tidySingleCellExperiment/commit/514b7f1ce20e58b1ae08844d8793d86234c59a8b |

## Methods

### Harmonisation and curation of the CELLxGENE atlas metadata

We downloaded the metadata and gene-transcript abundance from the CELLxGENE database (cellxgene.cziscience.com) through the R package cellxgenedp (10.18129/B9.bioc.cellxgenedp). We then harmonised and merged the metadata from 324 datasets, obtaining a dataset with consistent data types and ontologies. The abundance matrices were harmonised to have the same gene set, following the HUGO Gene Nomenclature^97^. We standardised the gene-transcript abundance distribution, defined in the real positive scale (e.g. not logged or square rooted). We standardised sample identifiers (named ‘sample_’ in our database) using a combination of study-specific identifiers and sample metadata. For example, cells grouped under the same identifiers but sourced from multiple tissues or individuals of distinct ages were split accordingly, as belonging to several samples by definition. We consistently represented age as days (named ‘age_days’ in our database). Where a categorical value was provided, we converted it into equivalent days using a publicly available reference^98^. For example, the “adolescent stage” was converted to 15 years old (= 5475 days). To increase the interpretability of results, we grouped 156 tissue labels (excluding nine cell cultures) in 45 tissue classes (named ‘tissue_harmonised’ in our database).

### Cell type annotation

Immune cells were labelled using Seurat Azimuth mapping to the PBMC reference^92^ and SingleR^16^ with the Blueprint^99^ and Monaco^100^ references^16^. To identify a consensus, we compared and contrasted the high-resolution labels ‘predicted.celltype.l2’ for Seurat Azimuth and ‘label.finè Blueprint and Monaco references. When possible, the reference-specific cell-type labels were standardised under a common ontology. Where the resolution of transcriptionally similar cell types was uncertain with the given tools, cell types were labelled with a coarser resolution. For example, innate lymphoid and natural killer cells were grouped under “innate lymphoid”. The cell type curation was performed to obtain a high confidence, meaningful representation of the immune system’s heterogeneity, allowing data-rich cell types whose tissue composition can be modelled probabilistically rather than aiming for the finest resolution possible.

The original annotation provided by the studies was integrated with the three new annotation sources to identify a absolute or relative consensus. We created confidence tiers based on how many annotation sources agreed. We gave a cell a confidence tier 1 if all four annotations agreed and a confidence tier of 2, 3, or 4 if 1, 2 or 3 annotations disagreed. Where not even a partial agreement across annotations was found, cells were labelled “other immune”. These cells were included in the compositional modelling but not tested for changes.

### Harmonised atlas query interface

To support data access and programmatic exploration of the harmonised atlas, we developed R and Python interfaces (*CuratedAtlasQuery*, Figure 2J). The query workflow is divided into exploration, filtering and data collection. The metadata is explorable in R using dbplyr^101^, which follows tidy principles^101^. The cell metadata is displayed in the form of a tidy table^101^. Cells represented in the table can be filtered for any condition based on the information relative to cells (e.g. cell type), samples (e.g. ethnicity), and datasets (e.g. tissue). The filtered table with cell identifiers can be used to download transcript abundance for the cells of interest. The transcript abundance is packaged as SingleCellExperiment (R), Seurat (R) or AnnData (Python) containers. When the container allows it, transcript abundance is saved into H5-based matrices, which can be analysed without overloading the machine’s memory^17^. Both the original transcript abundance and counts-per-million can be queried. Pseudobulk counts are also available to facilitate large-scale, summary analyses of transcriptional profiles. The metadata and transcript abundance matrices exist in the cloud and are hosted in the Object Store in the ARDC Nectar Research Cloud (for CuratedAtlasQuery version 0.99.1). The metadata is stored in the parquet format (parquet.apache.org).

### Data preparation and filtering for compositional analyses

The harmonised CELLxGENE atlas metadata was used for cellularity (proportion of immune cells among all cells) compositional analyses of immune cells. We selected primary (no re-analysed data or collections) physiological samples (i.e. no disease). We also excluded samples with less than 30 cells. We excluded erythrocytes and platelets from the analyses as they were not of interest. To increase the sample size per ethnicity group, we merged Asian descendants (labelled in CELLxGENE as Asian and Chinese). We excluded samples whose age was unknown (n=699) and scaled the age for the remaining samples (n=12282).

For immune cellularity analyses, we only considered non-blood and non-lymphoid tissues. We observed a bimodal distribution for the observed immune cellularity (arithmetically calculated). The modes of this distribution approach 0 (absence of immune cells) and 1 (absence of non-immune cells), where the valley of the distribution was at 0.75. Although our method is insensitive to outliers, we excluded samples that were experimentally enriched or depleted for immune cells (Supplementary File 2) with observed cellularity bigger than 0.75 to avoid large-scale biases. Similarly, to avoid samples that might have been experimentally depleted of immune cells, we selected those with one or more immune cells. For immune composition analyses, we filtered out non-immune cells. To ensure high-confidence analyses, we selected immune cells with confidence tier 1, 2 or 3.

For modelling immune cellularity, datasets with less than three samples were excluded. To model immune composition, we used more stringent filtering (compared with immune cellularity) for datasets with more than 15 samples, as the data availability (cell counts) for several cell types is lower than for the two immune and non-immune cell classes.

### Overview of the analysis design

In total, we produced eight models. Across the four factors: age, sex, ethnicity and technology, we modelled immune cellularity (proportion of immune cells among all cells) and composition. For each of these models, we test two main hypotheses, including the existence of a local (a.k.a. group or random) effect for each tissue and a global (a.k.a. population or fixed) effect at the body level. For the hypothesis testing of changes at the body level, just the population-level effect was considered. In contrast, the testing at the tissue level was done considering the sum of the population-level and group-level effects.

The reason for testing our hypotheses on distinct models (rather than testing all hypotheses on one only) is that distinct hypotheses might require distinct data subsets and intercepts (baseline) terms. For example, the age effect (as detailed below) is tested only in post-embryonic samples (> 1 year after conception); similarly, the sex effect is tested only for samples for which sex is known. Also, in visualising the data with the unwanted variation removed, it is helpful to choose an appropriate intercept (or no intercept) depending on the factors of interest. Additionally, reducing the model complexity for factors of no interest (which vary across hypotheses) is beneficial. This approach improves execution times and model convergence. For example, in testing for age, we do not need to resolve all cell-probing technologies; therefore, we can group the 10X chemistry versions, as the goal is to capture the variability at the technology level and not to estimate the effect for every chemistry. Following this rationale, the ethnicity label was simplified. The unknown or admixed ethnicities and those with less than 100 samples each were grouped within the label “other”, including Eskimo, Finnish, Pacific Islander, Oceanian, and Greater Middle Eastern. Similarly, unless the technology was a factor of interest, the 10X single-cell RNA sequencing chemistries and versions were grouped under the same class. Because of their unicity and this study focused on young-to-adult individuals, embryonic stages were excluded from the age, sex, and ethnicity-related analyses.

Unless stated otherwise, datasets (made unique if spanning more than one tissue) were modelled as group-level (a.k.a. random) intercept, and tissue was used as a local (a.k.a. group-level, random) effect for ageing. The global (a.k.a. population-level, fixed) effects were age, tissue, sex, ethnicity, and technology. The statistical inference was performed with a multi-level extension of sccomp (Supplementary Methods^19^) based on the Stan language^38^.

### Removal of unwanted variation

The removal of unwanted variation is based on the calculation of residuals^102^. The estimates are first produced for the model of choice. A target model is specified for the effects of interest. Then, the residuals are calculated between the observed data and the data generated from the target model. For example, if the fitted model is *∼ ‘factor of interest’ + ‘unwanted variation’ + (1 + ‘factor of interest’ | grouping)*, and the effects to preserve are defined as *∼ ‘factor of interest’*, data will be generated from the latter model excluding hierarchical and non-hierarchical unwanted variability. If the effects of interest include random effects, the desired formula is *∼ ‘factor of interest’ + (1 + ‘factor of interest’ | grouping)*.

### Modelling age effects

To model age association with immune cellularity (proportion of immune cells among all cells) and composition, samples less than one year after conception were excluded as the analysis focused on old-age ageing rather than early development. The variability was modelled to be associated with age and tissue. To visualise the trend of immune cellularity in ageing, we removed all variations in the estimated abundance and variability, except for age and the population effect.

### Modelling sex effects

To model sex association with immune cellularity (proportion of immune cells among all cells) and composition, samples with unknown sex were excluded. Tissues with less than one represented sex were excluded. To increase the signal-to-noise ratio, as sex differences are most observed from adolescence, samples less than 15 years old were excluded. The model was set with females as intercept (baseline). The variability was modelled to be associated with sex and tissue. To visualise the trend of immune cellularity between sexes, we removed all variation in the estimated abundance and variability, except for the sex and the population effect.

### Modelling ethnicity effects

To model ethnicity association with immune cellularity (proportion of immune cells among all cells) and composition, samples with unknown ethnicity were excluded. The model was set to intercept free. The variability was modelled to be associated with ethnicity and tissue. To visualise the trend of immune cellularity between sexes, we removed all variation in the estimated abundance and variability, except for the sex and the population effect.

### Modelling the tissue immune landscape

To create a reference map of immune cellularity and composition across tissues, we estimated those quantities after accounting for unwanted biological (i.e. age, sex and ethnicity) and technical (i.e. technology and dataset) variability. The model was set to intercept free. Datasets were used to estimate group-level tissue effects. The estimated cellularity and immune composition per tissue were calculated by factoring out technology and group-level dataset effects. The estimated cellularity was converted to proportion with an inverse softmax transformation. The dimensionality reduction was performed on the original data, and the data with unwanted variation was removed, as described in the Methods subsection (Removal of unwanted variation). Dimensionality reduction was performed with tidybulk^20^.

### Cell communication analyses

We tested changes in communication patterns in ageing across tissues. To ensure high-confidence analyses, we selected immune cells with confidence tier 1, 2 or 3. After filtering, for each sample, we excluded cell types with less than ten cells for each sample. Then, we excluded samples with less than two remaining immune cell types. The scoring of cell communication was done with CellChat^37^ with default parameters. For each sample, we calculated the communication weight across communication axes. We included cell-cell interaction, extracellular-matrix interaction (for solid tissues), and secreted factors. Two weights were calculated as the overall communication strength of a cell type to all other cell types and the communication strength between each cell type pair. To ensure quality results, as the inference of cell communication adds uncertainty to the hypothesis testing, we limited our modelling to the tissues with a large sample size after filtering, for which communication was detected. We included blood, lymph nodes, heart and lung. To account for group-level (a.k.a. random) effects, we used brms for multilevel statistical modelling^13^. We used a beta noise model, given that the weight was bounded between 0 and 1.

### Tissue-residency analyses

We scored tissue residency for T cells using Singscore^93^ and the residency reference Mckay et al.^33^, on the scaled counts (counts per million). For comparative purposes, we estimated the residency score using the signature Bhuva et al.^93^ proposed, which showed a high correlation (Supplementary Figure S5). As residency and exhaustion are related phenomena, we used Singscore to score T cell exhaustion. However, exhaustion was not included in the analyses as no cell showed a higher-than-background score. (Supplementary Figure S6).

### Differential gene transcript abundance analyses

We used the glmmSeq algorithm to perform multilevel (random effect) transcriptional analyses^12^ implemented in tidybulk^20^, including 116,606 pseudobulk samples, resolved for cell types. Two types of analyses were performed. In the tissue-grouped analyses, we split the dataset across tissues, and the cell types were used as a grouping in the multilevel (i.e. random effect) models. In the cell-type-grouped analyses, we split the dataset across cell types, and the tissues were used as a grouping in the multilevel (i.e. random effect) models. The tissue-grouped analyses would identify genes where transcription was altered across cell types for each tissue. In contrast, the cell-type-grouped analyses would identify genes where transcription was altered across tissue for each cell type. GlmmSeq was used with the lme4 backend, where the feature-wise dispersion was calculated using edgeR^103^. If more than 2000 pseudobulks were present in a dataset (e.g., in blood tissue), the glmmmTMB backend was used instead. To parallelise the analyses, we used the workflow manager R Targets (CRAN repository). Transcriptional analyses were performed at the pseudobulk level resolved for samples and cell types. The pseudobulk counts were calculated with tidyseurat^104^. Considering the diversity of standards in the studies included in *CuratedAtlasQuery*, counts were normalised using quantile normalisation using tidybulk^20^. Lowly abundant gene transcripts were excluded, using edgeR^103^ through tidybulk (minimum_counts = 500, minimum_proportion = 0.9) and using sex and ethnicity as factors of interest. Two models were used to estimate transcriptional abundance, depending on whether the analyses were carried out across cell types or tissues. For cell types, we estimated the following model for each cell type ∼ age * sex + ethnicity + technology + number_aggregated_cells + disease + (1 + age * sex + ethnicity | tissue). For tissues, we estimated the following model for each tissue ∼ age * sex + ethnicity + technology + number_aggregated_cells + disease + (1 + age * sex + ethnicity | cell_type). For each estimation, we excluded the samples that caused complete confounding of the covariates; also, we excluded the covariates that did not have enough diversity (e.g. sex where females were not present). In the tissue-grouped analyses, tissues were excluded if not more than three cell types were present. In the cell type-grouped analyses, cell types were excluded if no more than three tissues were present.

### Ageing trajectory analyses

To predict compositional ageing program and tissue trajectories, we used sccomp (sccomp_predict) to predict the body-level and tissue-level effects of ageing on composition with, encoded in the formula ‘composition ∼ age + (age | tissue)’. We predicted the tissue immune composition for ages from 10 to 90 years with intervals of 10 years. From such prediction, we transform the proportion to a logit scale and calculate the PCA (using prcomp) without scaling.

### Software for data manipulation and visualisation

The data analysis was performed in R^105^. Data wrangling was done through tidyverse^101^. Single-cell data analysis and manipulation were done through Seurat^95^ (version 4.0.1), tidyseurat^104^ (version 0.3.0), tidySingleCellExperiment^96^ (version 1.9.4), and tidybulk^20^ (version 1.6.1). Colour-blind-friendly colouring was aided by dittoseq^106^. Heatmaps were created with ggplot2^107^ and tidyHeatmap^108,109^. GGupset was used for upset plots. Parallelisation was achieved through makeflow^110^.

