## Supplementary Methods and Figures for "A multi-organ map of the human immune system across age, sex and ethnicity"

### Supplementary Methods

#### *Atlas-level compositional and variability analysis*

The core model[1] allows compositional and variability analyses that are robust against outliers. Motivated by the complex and hierarchical structure of the CELLxGENE and other single-cell atlases, we extended this method to model hierarchical effects (Figure 1B), including tissue and dataset. The user can express the hierarchy structure analogously to lme4[2]. For example, we can use tissue as a grouping variable to estimate a global effect (e.g. age) throughout the body. In this scenario, we would observe a significant effect if the effect is observed in the vast majority of tissue (rather than the majority of samples). This multilevel structure is essential in large-scale atlas-level analyses. Otherwise, the estimates would be biased by the most represented tissues (e.g. blood). Hierarchical effects can also be captured at the dataset level, for example, on the intercept.

Our method can remove unwanted group- and population-level effects for arbitrary linear models. We achieve this with a residual-based strategy (Methods), in which the deviance from the mean effect is calculated from the joint-posterior distribution. Uniquely, our method can calculate the uncertainty of the adjusted data based on the uncertainty of the factors of interest. This ability is important for deconvolving and visualising effects of interest from complex data.

#### *Multilevel modelling*

Multilevel modelling suits complex designs where observations are hierarchically dependent, similar to students in schools. We implemented the multilevel modelling extending the core model[1]. The multilevel modelling is expressed in the formula in style used in the lme4[2] and brms[3] R packages. We separate the multilevel model into the population effects (also called fixed effects) and the group effects (also called random effects) (Equation 1).

$$M = \Gamma X + \Psi Z \qquad (1)$$

***Γ*** and ***Ψ*** are population- and group-level coefficients, respectively, and ***X*** and ***Z*** are the corresponding design matrices. ***M*** is a matrix with dimensions G×S (from [1]) where G is the number of cell groups, and S is the number of samples. ***Γ*** is a matrix with dimensions G×C where C is the number of population-level coefficients. ***Ψ*** is a matrix with dimensions G×R, where R is the number of groups. ***X*** is a matrix with dimensions C×S, and ***Z*** has dimensions R×S. To model the correlation structure among the random effects, we impose a multinormal prior to ***Ψb*** shared across cell types (Equation 3). This prior is centred at zero, with the covariance matrix Ω of size C (dimension C square), describing the variability and correlation structure. The covariance matrix Ω is created by defining a standard deviation component σ of size C and a correlation matrix of size C (Equation 4). The Correlation matrix and variance are given non-informative priors (Equations 5 and 6).

$$\Psi_g \sim multinormal(0, \Omega) \qquad (3)$$

$$\Omega_C = f(\sigma_C, \Upsilon_C) \qquad (4)$$

$$\Upsilon \sim LewandowskiKurowickaJoe(2) \qquad (5)$$

$$\sigma \sim LogNormal(0, 1) \qquad (6)$$

### Supplementary Tables and Figures

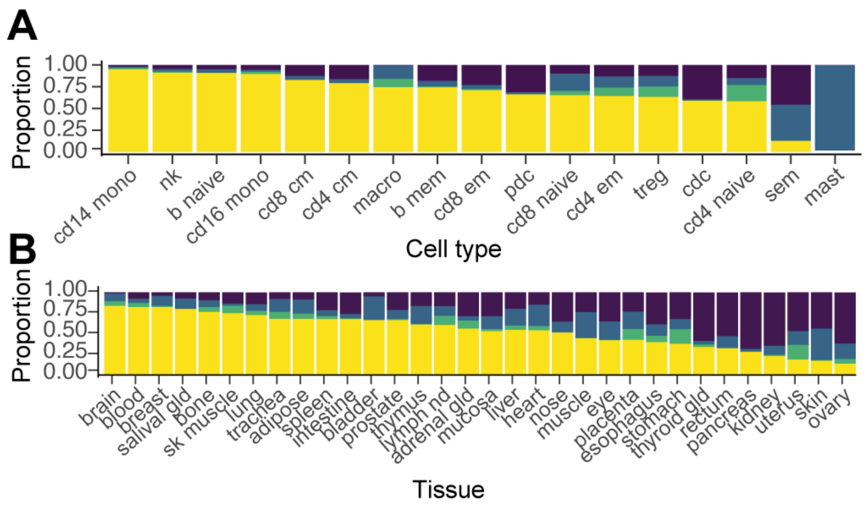


**Supplementary Figure S1. A:** Proportion of confidence classes across cell types. Cell types have a variable confidence tier distribution, with immune progenitor cells and mast cells having the worst confidence. **B:** Proportion of confidence classes across tissues. For example, the brain, blood and breast have nearly 100% high-confidence tiers, while the ovary, kidney, and pancreas have only 25% high-confidence tiers.

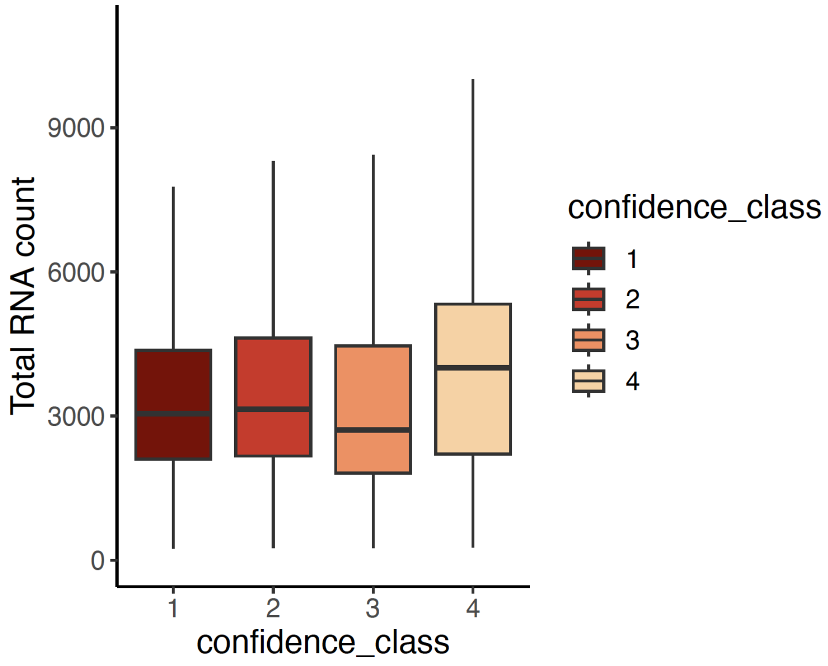


**Supplementary Figure S2.** Association between confidence tier (x-axis) and library size (y-axis) for the samples with unscaled integer counts expressing transcriptional abundance.

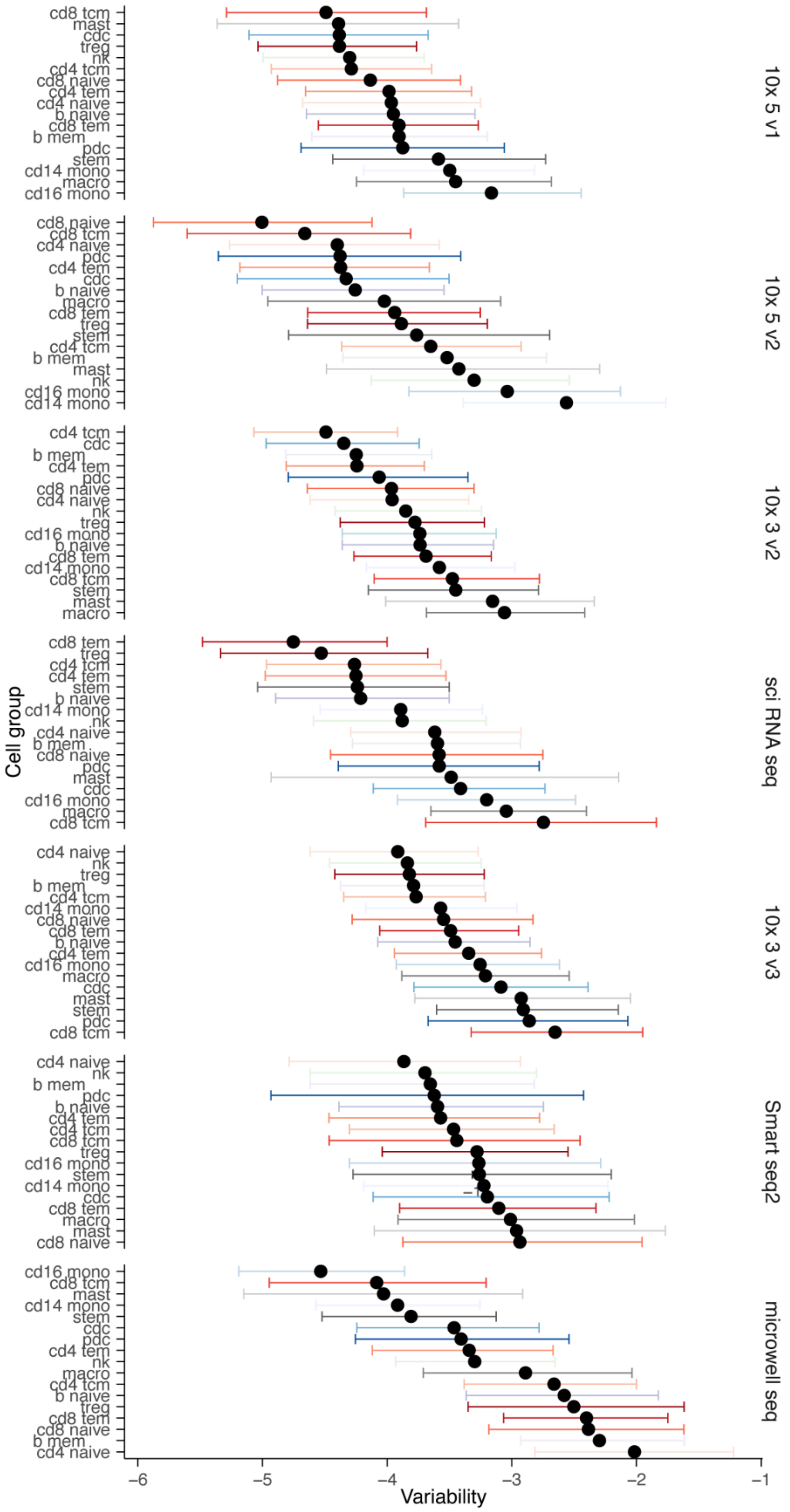
10x 5 v1
cd8 tcm
mast
cdc
treg
nk
cd4 tcm
cd8 naive
cd4 tem
cd4 naive
b naive
cd8 tem
b mem
pdc
stem
cd14 mono
macro
cd16 mono
10x 5 v2
cd8 naive
cd8 tcm
cd4 naive
pdc
cd4 tem
cdc
b naive
macro
cd8 tem
treg
stem
cd4 tcm
b mem
mast
nk
cd16 mono
cd14 mono
10x 3 v2
cd4 tcm
cdc
b mem
cd4 tem
pdc
cd8 naive
cd4 naive
nk
treg
cd16 mono
b naive
cd8 tem
cd14 mono
cd8 tcm
stem
mast
macro
sci RNA seq
cd8 tem
treg
cd4 tcm
cd4 tem
stem
b naive
cd14 mono
nk
cd4 naive
b mem
cd8 naive
pdc
mast
cdc
cd16 mono
macro
cd8 tcm
10x 3 v3
cd4 naive
nk
treg
b mem
cd4 tcm
cd14 mono
cd8 naive
cd8 tem
b naive
cd4 tem
cd16 mono
macro
cdc
mast
stem
pdc
cd8 tcm
Smart seq2
cd4 naive
nk
b mem
pdc
b naive
cd4 tem
cd4 tcm
cd8 tcm
treg
cd16 mono
stem
cd14 mono
cdc
cd8 tem
macro
mast
cd8 naive
microwell seq
cd16 mono
cd8 tcm
mast
cd14 mono
stem
cdc
pdc
cd4 tem
nk
macro
cd4 tcm
b naive
treg
cd8 tem
cd8 naive
b mem
cd4 naive
Cell group
−6
−5
−4
−3
−2
−1
Variability

**Supplementary Figure S3.** Schematics of the statistical modelling. On the left-hand side, we show input counts and the hierarchically organised metadata. In the middle-top, we show the outlier identification layer and the hierarchical fit, with adaptive shrinkage of the variance given by the information sharing across cell types. In the middle-bottom, we show the parametrisation of the population-level (a.k.a. fixed) and group-level (a.k.a. random) effects. On the right-hand side, we show the hypothesis testing and removal of unwanted variation modules.

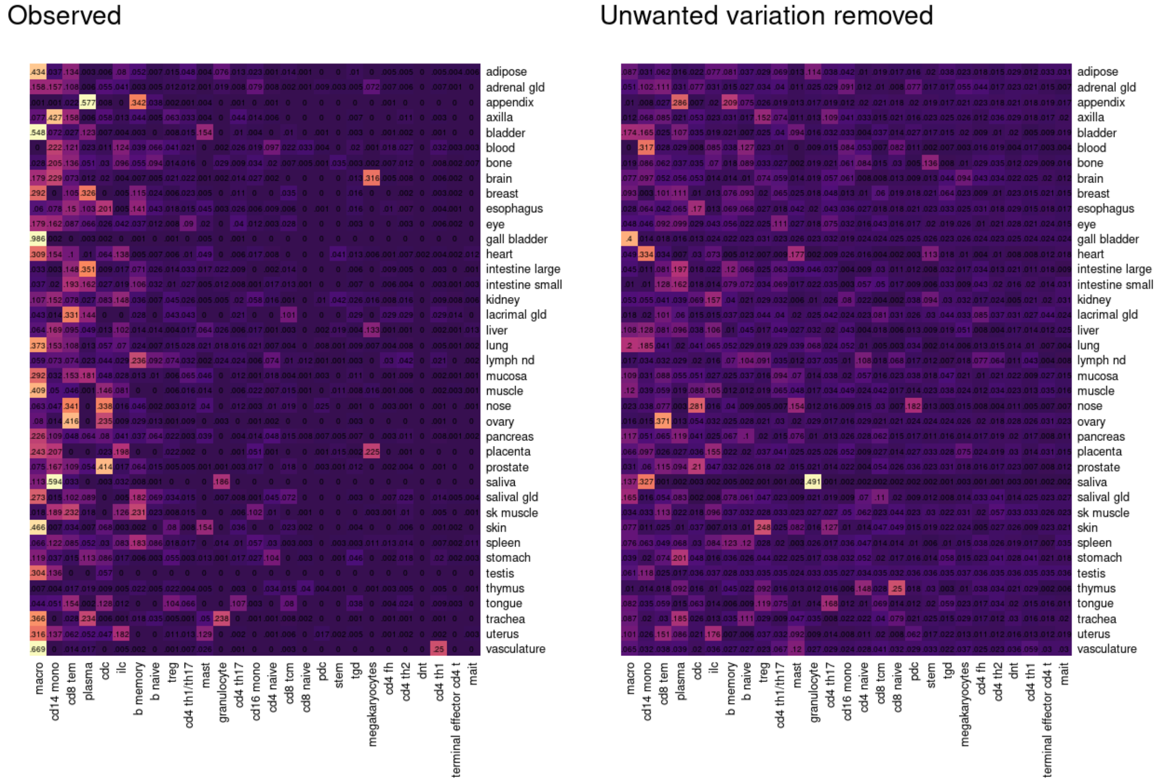

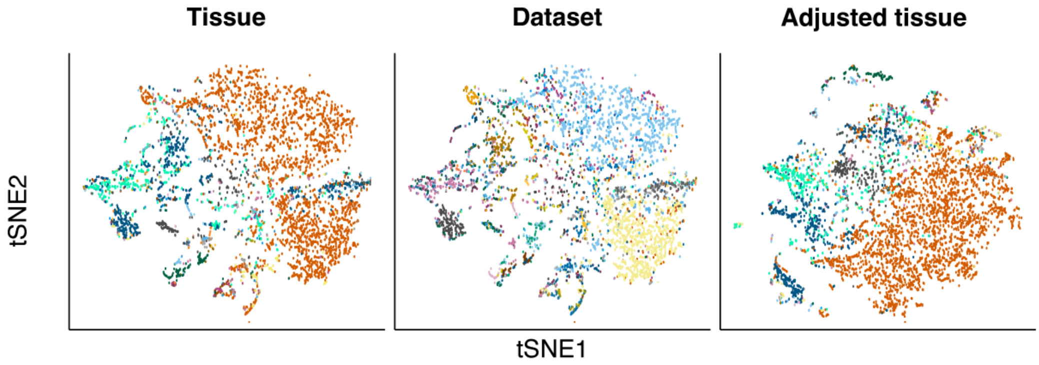


**Supplementary Figure S4. Top:** Comparison of the reference map derived from the arithmetic calculation of cell-type proportion (which includes confounders such as age, sex, ethnicity, dataset, and probing technology) and the estimated mean proportion with the unwanted variation removed. **Bottom:** Distribution of samples in tSNE space according to immune composition. Data points are coloured by tissues/tissue in the left and right panels and by datasets in the middle. Adjusted composition (right panel) refers to removing unwanted biological and technical variation as specified in the statistical model (grey box, Figure 4).

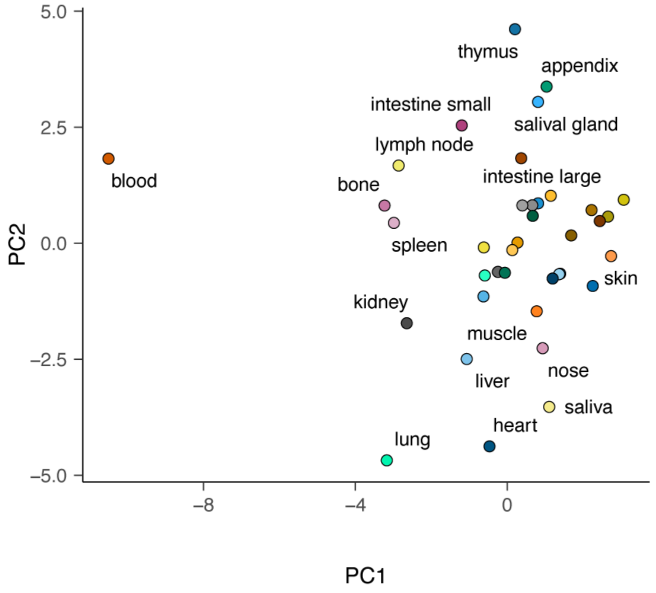

**Figure S5.** Distribution of tissues based on their similarity in immune makeup (including all immune cells tyme and non-immune cells), excluding technical and biological effects of age, sex, ethnicity, and technology.

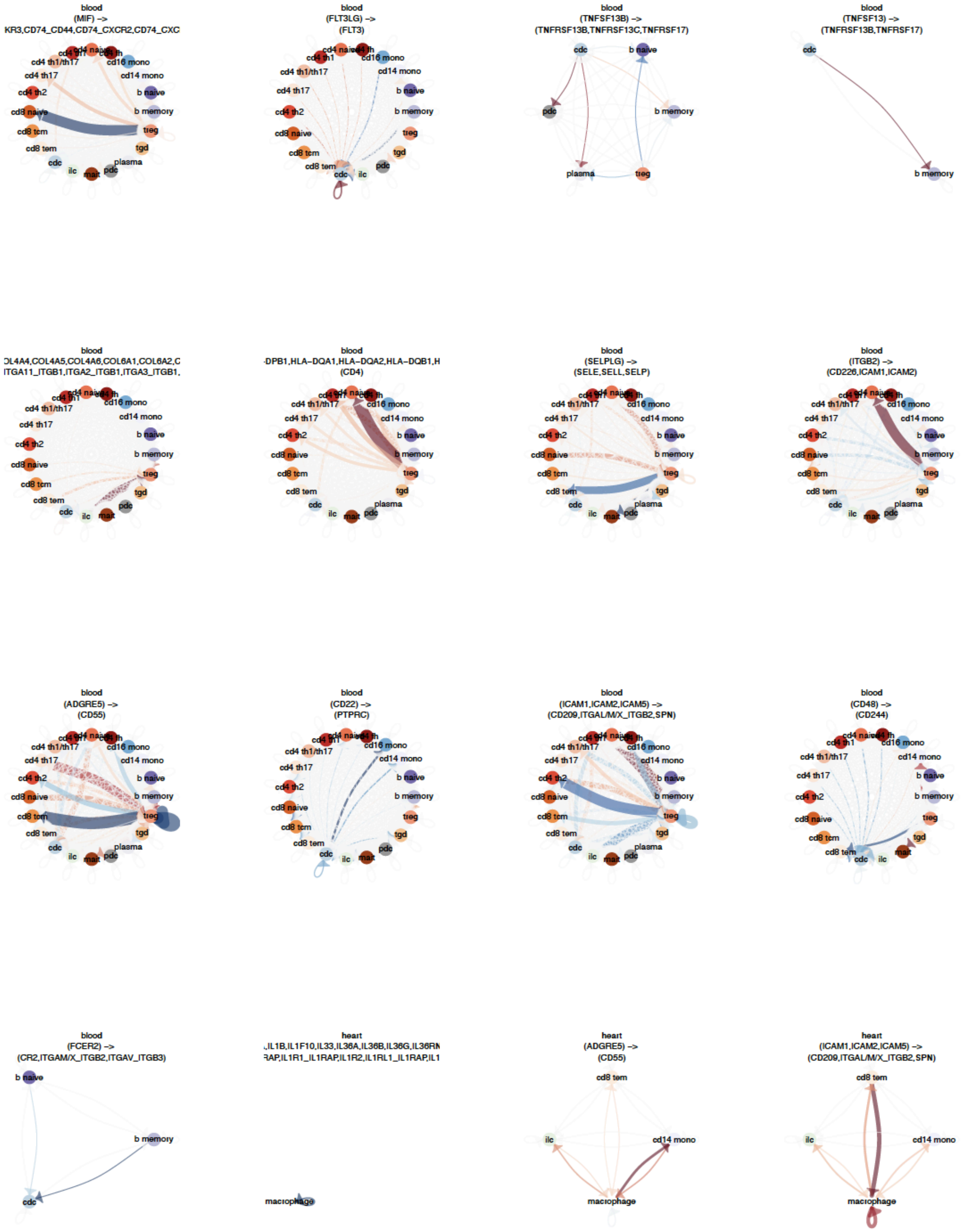
blood
(MIF) ->
KR3,CD74_CD44,CD74_CXCR2,CD74_CXC
blood
(FLT3LG) ->
(FLT3)
blood
(TNFSF13B) ->
(TNFRSF13B,TNFRSF13C,TNFRSF17)
blood
(TNFSF13) ->
(TNFRSF13B,TNFRSF17)
blood
OL4A4,COL4A5,COL4A6,COL6A1,COL6A2,C
ITGA11_ITGB1,ITGA2_ITGB1,ITGA3_ITGB1,
blood
-DPB1,HLA-DQA1,HLA-DQA2,HLA-DQB1,H
(CD4)
blood
(SELPLG) ->
(SELE,SELL,SELP)
blood
(ITGB2) ->
(CD226,ICAM1,ICAM2)
blood
(ADGRE5) ->
(CD55)
blood
(CD22) ->
(PTPRC)
blood
(ICAM1,ICAM2,ICAM5) ->
(CD209,ITGAL/M/X_ITGB2,SPN)
blood
(CD48) ->
(CD244)
blood
(FCER2) ->
(CR2,ITGAM/X_ITGB2,ITGAV_ITGB3)
heart
,IL1B,IL1F10,IL33,IL36A,IL36B,IL36G,IL36RN
RAP,IL1R1_IL1RAP,IL1R2,IL1RL1_IL1RAP,IL1
heart
(ADGRE5) ->
(CD55)
heart
(ICAM1,ICAM2,ICAM5) ->
(CD209,ITGAL/M/X_ITGB2,SPN)

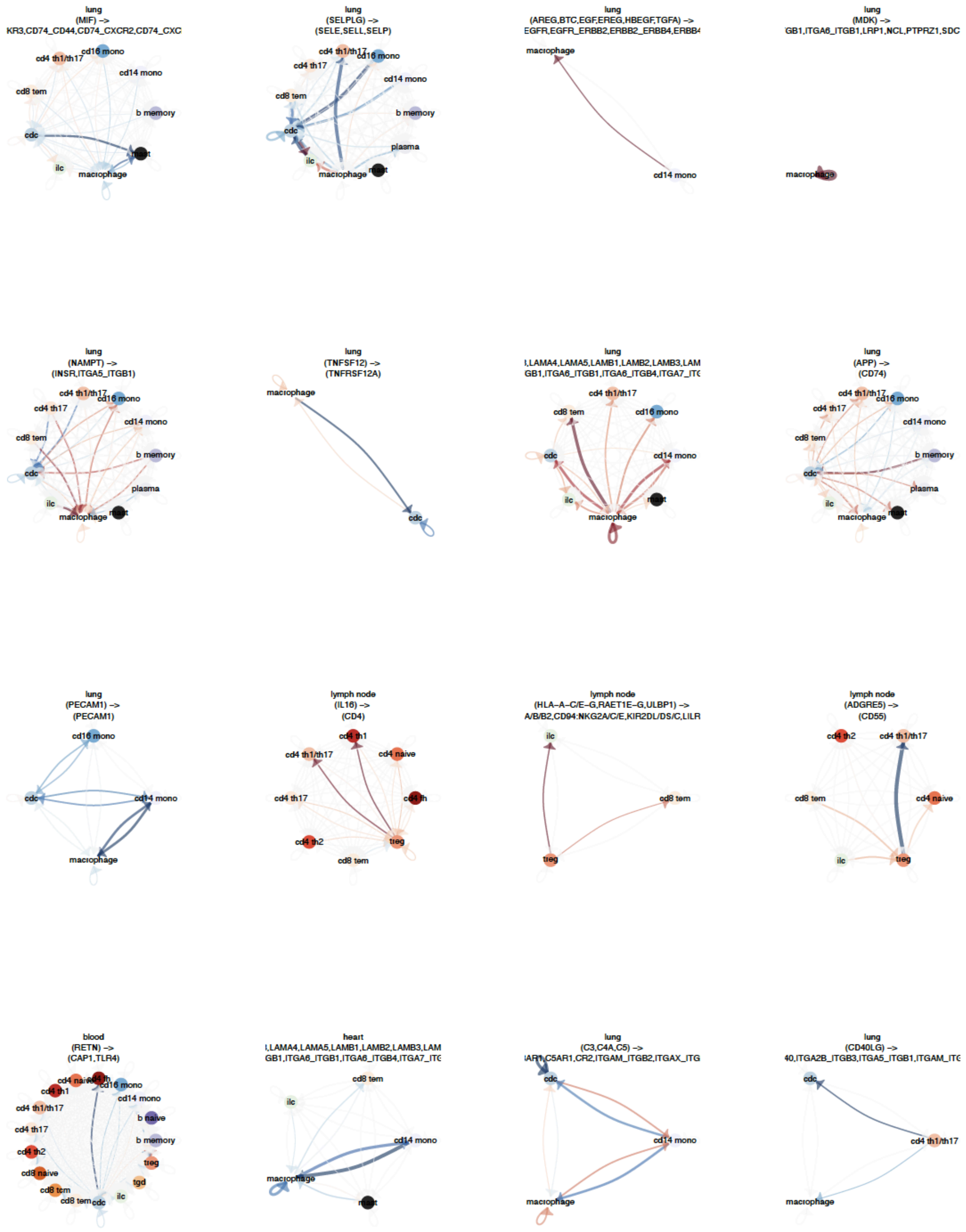
lung
(MIF) ->
KR3,CD74_CD44,CD74_CXCR2,CD74_CXC
cd16 mono
cd4 th1/th17
cd14 mono
cd8 tem
b memory
cdc
mast
ilc
macrophage
lung
(SELPLG) ->
(SELE,SELL,SELP)
cd4 th1/th17
cd16 mono
cd4 th17
cd14 mono
cd8 tem
b memory
cdc
plasma
ilc
macrophage
mast
lung
(AREG,BTC,EGF,EREG,HBEGF,TGFA) ->
EGFR,EGFR_ERBB2,ERBB2_ERBB4,ERBB4
macrophage
cd14 mono
lung
(MDK) ->
GB1,ITGA6_ITGB1,LRP1,NCL,PTPRZ1,SDC
macrophage
lung
(NAMPT) ->
(INSR,ITGA5_ITGB1)
cd4 th1/th17
cd16 mono
cd4 th17
cd14 mono
cd8 tem
b memory
cdc
plasma
ilc
macrophage
mast
lung
(TNFSF12) ->
(TNFRSF12A)
macrophage
cdc
lung
,LAMA4,LAMA5,LAMB1,LAMB2,LAMB3,LAM
GB1,ITGA6_ITGB1,ITGA6_ITGB4,ITGA7_ITG
cd4 th1/th17
cd8 tem
cd16 mono
cdc
cd14 mono
ilc
mast
macrophage
lung
(APP) ->
(CD74)
cd4 th1/th17
cd16 mono
cd4 th17
cd14 mono
cd8 tem
b memory
cdc
plasma
ilc
macrophage
mast
lung
(PECAM1) ->
(PECAM1)
cd16 mono
cdc
cd14 mono
macrophage
lymph node
(IL16) ->
(CD4)
cd4 th1
cd4 th1/th17
cd4 naive
cd4 th17
cd4 tfh
cd4 th2
treg
cd8 tem
lymph node
(HLA-A-C/E-G,RAET1E-G,ULBP1) ->
A/B/B2,CD94:NKG2A/C/E,KIR2DL/DS/C,LILR
ilc
cd8 tem
treg
lymph node
(ADGRE5) ->
(CD55)
cd4 th2
cd4 th1/th17
cd8 tem
cd4 naive
ilc
treg
blood
(RETN) ->
(CAP1,TLR4)
cd4 naive
cd16 mono
cd4 th1
cd14 mono
cd4 th1/th17
b naive
cd4 th17
b memory
cd4 th2
treg
cd8 naive
tgd
cd8 tcm
cd8 tem
cdc
ilc
heart
,LAMA4,LAMA5,LAMB1,LAMB2,LAMB3,LAM
GB1,ITGA6_ITGB1,ITGA6_ITGB4,ITGA7_ITG
cd8 tem
ilc
cd14 mono
macrophage
mast
lung
(C3,C4A,C5) ->
AR1,C5AR1,CR2,ITGAM_ITGB2,ITGAX_ITG
cdc
cd14 mono
macrophage
lung
(CD40LG) ->
40,ITGA2B_ITGB3,ITGA5_ITGB1,ITGAM_ITG
cdc
cd4 th1/th17
macrophage

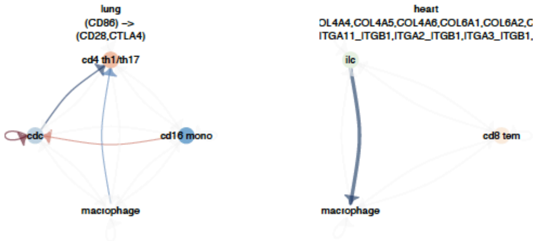


**Supplementary Figure S6.** Communication graphs for the significantly altered axes of communication between T regulatory cells, monocytes or dendritic cells and any other cell type. Blue and red colours represent downregulation and upregulation, respectively. The list of genes of each communication axis is within brackets.

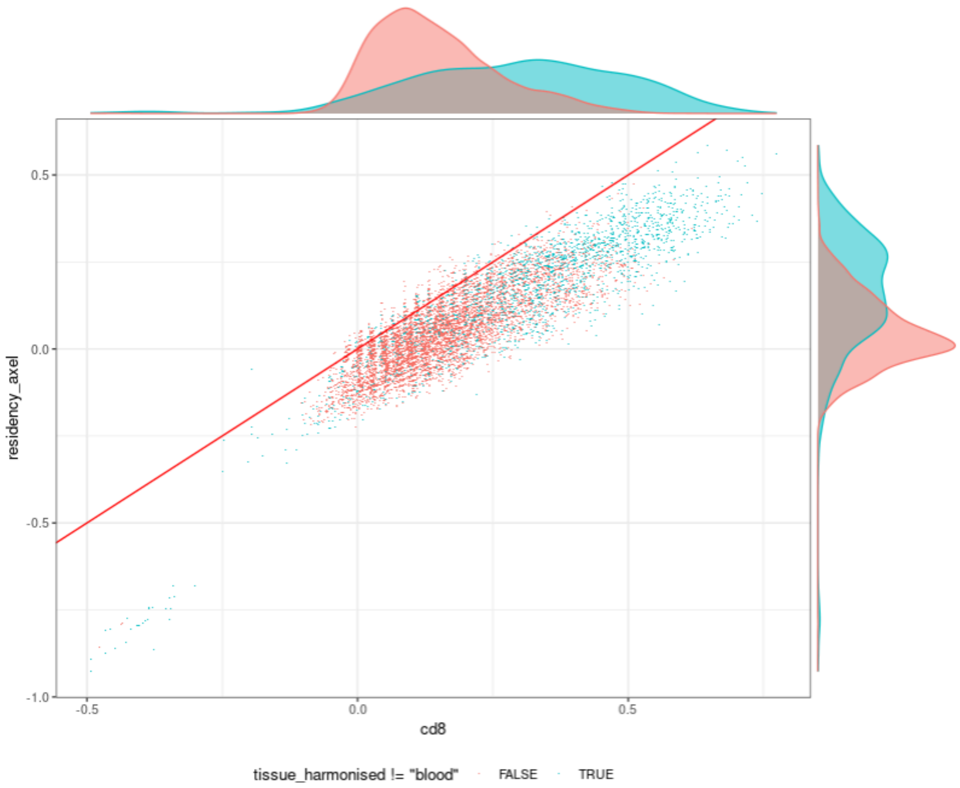


**Supplementary Figure S7.** Correlation between the residency signature from the Mackay 2016 residency signature (DOI: 10.1126/science.aad203) and the Foroutan 2021 residency signature (DOI: 10.1158/2326-6066.CIR-21-0137).

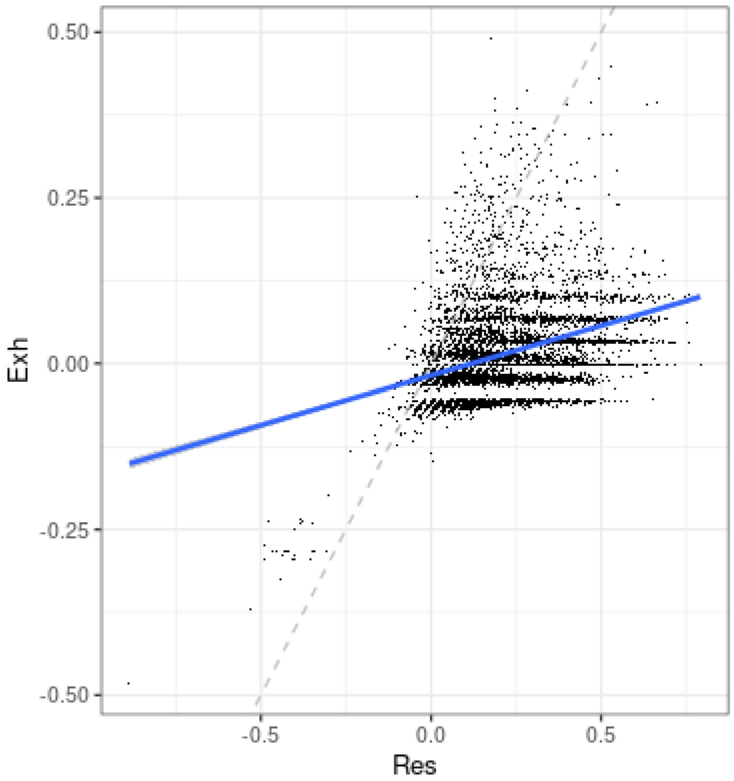


**Supplementary Figure S8.** Correlation between the residency and exhaustion signature from Foroutan et al. 2021 (DOI: 10.1158/2326-6066.CIR-21-0137). A value higher than 0.25 represents a detectable signature.

### Supplementary Files

**SUPPLEMENTARY_datasets_to_check_in_the_literature_checked.csv**

This file includes the curation of the datasets used for cellularity estimation. We checked whether the datasets were enriched or depleted for immune cells, and/or specific immune cell types.

**SUPPLEMENTARY_estimates.xlsx**

This file includes the statistics for the cellularity and compositional analyses performed in this study.

### SUPPLEMENTARY RESULTS

#### Removal of technology effect from the measurement of tissue composition

Given the diversity of this resource, we next sought to explore the technologies and protocols used to probe cell RNA and identify technical effects in cell type representation across tissues (Supplementary File 1).

The median number of cells assayed in each sample differs between technologies, ranging from <10 cells by Patch-Seq to 19,411 by Slide-Seq (Figure 4A, top). The atlas incorporated 19 distinct protocols (Figure 4A, bottom). Nine used 10X microfluidics, representing 70.2% of all cells. Other highly represented technologies include sci-RNA-seq[4] and Microwell-seq. A total of 814 tissue specimens were assayed with multiple technologies (10X and Smart-Seq2[5]; Figure 4B), making this atlas a powerful resource for technology benchmarking.

Principal component analysis of the cellular composition, adjusted for unwanted variation, shows that the 10x technologies were consistent and did not show significant compositional differences. In contrast, Smart-seq2, microwell-seq[6] and sci-seq showed significant differences along the second principal components (Figure 4C). These differences highlight the importance of including the technology in the model to improve compositional estimates. We then investigated which cell types mainly contributed to the technology effect.

Compared to the most common technology, 10x 3' version-2, we detected several significant differences with other assays (Figure 4D). The number of significant differences per technology reflected the distances observed in the principal component plot. Sci-seq and microwell-seq showed the most significant differences with 9 and 7, respectively. The set of cell types having significant differences between these two methods did not show considerable overlap. Despite Smart-seq2 being a fundamentally different technology to 10x, it did not show significant differences at the cell-type level.

Consistency in probing cell types of interest should be considered when choosing the technological platform. Therefore, we tested the differential variability of cellularity composition across tissues (Figure 4E, Supplementary Figure S3). 10x technologies were the most homogeneous (low variability), independent of other biological batch factors (Figure 3, grey box). Similarly, Sci-seq[7] showed low variability, although the uncertainty around the variability estimate is high due to the more limited data availability. Smart-seq2 and microwell-seq show the biggest compositional variability.

Modelling and excluding the technology effect on tissue composition allows to better resolve the major biological effects on the immune system across the human population, including age, sex, and ethnicity.

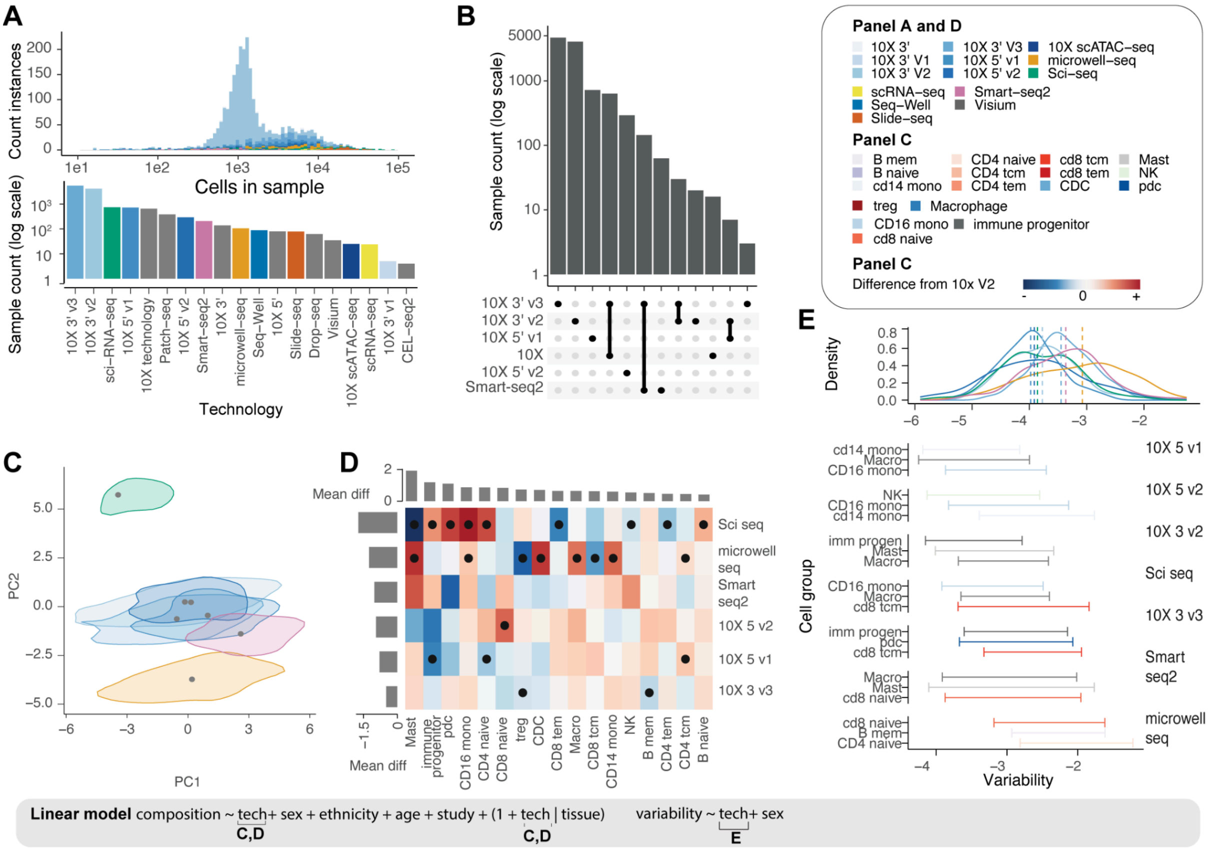


**Supplementary Figure S9. Unwanted variation in composition due to technology. A:** Distribution of the number of cells in each sample, coloured by technology (top) and the number of samples by technology (bottom). **B:** Counts of samples across multiple technologies. **C:** Principal component analysis of the immune system composition for the seven most represented technologies across tissues, independent of datasets, sex, age, and ethnicity (linear model described in the legend). Dots represent the expected dimensional value. The density areas represent the uncertainty around the expected value (density 0.02). **D:** Technology effects on cell type relative abundance, with the popular 10X 3’ v2 as a baseline. The bar plots represent the mean significant absolute change across technologies (rows) and cell types (columns). The dots represent the significant differences (false-discovery rate < 0.05). **E:** Consistency (i.e. differential variability) in cell type relative abundance across technologies independently from datasets, sex, age, ethnicity and tissue. The top panel represents the mean variability across cell types. The dashed lines represent the expected value, while the density lines represent the uncertainty. The bottom panel shows the variability 95% credible interval for the three least consistent (most variable) cell types per technology.
